# Intersecting impact of CAG repeat and Huntingtin knockout in stem cell-derived cortical neurons

**DOI:** 10.1101/2025.02.24.639958

**Authors:** Jennifer T. Stocksdale, Matthew J. Leventhal, Stephanie Lam, Yu-Xin Xu, Yang Oliver Wang, Keona Q. Wang, Reuben Tomas, Zohreh Faghihmonzavi, Yogi Raghav, Charlene Smith, Jie Wu, Ricardo Miramontes, Kanchan Sarda, Heather Johnson, Min-Gyoung Shin, Terry Huang, Mikelle Foster, Mariya Barch, Naufa Armani, Chris Paiz, Lindsay Easter, Erse Duderstadt, Vineet Vaibhav, Niveda Sundararaman, Dan P. Felsenfeld, Thomas F. Vogt, Jennifer Van Eyk, Steve Finkbeiner, Julia A. Kaye, Ernest Fraenkel, Leslie M. Thompson

## Abstract

Huntington’s Disease (HD) is caused by a CAG repeat expansion in the gene encoding Huntingtin (HTT). While normal HTT function appears impacted by the mutation, the specific pathways unique to CAG repeat expansion versus loss of normal function are unclear. To understand the impact of the CAG repeat expansion, we evaluated biological signatures of HTT knockout (*HTT* KO) versus those that occur from the CAG repeat expansion by applying multi-omics, live cell imaging, survival analysis and a novel feature-based pipeline to study cortical neurons (eCNs) derived from an isogenic human embryonic stem cell series (RUES2). *HTT* KO and the CAG repeat expansion influence developmental trajectories of eCNs, with opposing effects on the growth. Network analyses of differentially expressed genes and proteins associated with enriched epigenetic motifs identified subnetworks common to CAG repeat expansion and *HTT* KO that include neuronal differentiation, cell cycle regulation, and mechanisms related to transcriptional repression and may represent gain-of-function mechanisms that cannot be explained by *HTT* loss of function alone. A combination of dominant and loss-of-function mechanisms are likely involved in the aberrant neurodevelopmental and neurodegenerative features of HD that can help inform therapeutic strategies.

## Introduction

Huntington’s disease (HD) is an autosomal dominant brain disorder that affects approximately 10 in every 100,000 individuals^1^. HD typically manifests clinically at midlife; however, juvenile HD cases also occur and display more severe clinical manifestation and faster disease progression^2^. HD is caused by an unstable CAG repeat expansion in exon 1 of the *Huntingtin* (*HTT*) gene^3^. The resulting mutated gene encodes an elongated polyglutamine repeat (polyQ)-containing HTT protein. The length of the repeat expansion is highly polymorphic and in general correlates with disease severity: individuals with fewer than 36 repeats are unaffected, those with 36–39 repeats have variable penetrance^4^, adult-onset HD occurs in individuals with 40-60 repeats, and juvenile-onset HD typically occurs in individuals with more than 60 repeats^5^. The hallmark clinical features of HD are the progressive loss of physical and mental functions characterized by uncontrolled movements, and psychiatric and cognitive impairment^1^. The mutation causes dysfunction and death of medium spiny neurons in the striatum and significant dysfunction and atrophy of the cortex. During the past three decades, great progress has been made in understanding mechanisms involved in HD pathogenesis^6,7^, which have primarily centered around a dominant gain of aberrant function, and in identifying genetic modifiers of disease onset, which to date are largely functionally related to genome stability and DNA mismatch repair^8,9^.

There has also been significant progress in understanding normal HTT function, potentially as a protein scaffold for processes such as autophagy^10,11^, vesicular transport^12^, neurodevelopment^13,14^, and a broad range of other cellular processes (reviews^15–17^). While loss of HTT function is not sufficient to cause HD, given that genetic mutation or deletion of *HTT* alleles in human disorders including Wolf-Hirschhorn Syndrome^18^ do not cause a neurodegenerative disease similar to HD, the CAG repeat expansion does cause an apparent disruption of normal HTT functions including vesicular transport of BDNF^19^, mitotic spindle formation^20^, and autophagosome formation^21^. Further, homozygous CAG repeat expansion does not result in consistently worse disease than a heterozygous mutation in human HD^22,23^, reflecting the dominant nature of the mutation, whereas total loss of HTT is embryonic lethal in mice^24^. Some phenotypes impacted by expanded repeat HTT and HTT knockout (KO) are overlapping, including those impacting neurodevelopment and chromosomal stability^20^. However, to date, there has not been a systematic investigation of the degree to which HD-associated neuronal molecular and morphological changes are due to CAG repeat expansion versus loss of HTT function.

To address this, isogenic embryonic stem cell (ESC) lines with normal, CAG-expanded and a homozygous deletion of *HTT* exon 1, resulting in loss of the HTT protein expression (*HTT* KO), were differentiated to generate cortical-like neurons (eCNs) and a comprehensive analysis using live cell imaging and multi-faceted OMICs was conducted.

Previous time-lapse imaging using robotic microscopy (RM) of live neurons patterned from induced pluripotent stem cells (i-neurons) from HD patients and controls demonstrated neurodegenerative phenotypes, including increased cell death, greater susceptibility to trophic factor withdrawal, and differences in neurite length^25,26^. However, variability between patient-derived lines has been well-documented^27,28^ and previously observed phenotypes may have based on genetic or confounding factors unrelated to HTT pathology. Here, we report a subset of similar findings as previous studies in HD i-neurons^25,26^, but in isogenic CAG-expanded lines, suggesting that these cellular changes are directly attributable to the CAG repeat expansion. Interestingly, we do not observe similar changes in *HTT* KO lines, suggesting that these phenotypes are specifically due to a gain-of-function mechanism driven by the polyglutamine expansion, a mechanism not previously recognized.

We also developed a novel platform for image-based feature classification, commonly used in machine learning (ML) applications^29^, as a way to assess dynamic morphologic changes over time. Previously, ML called a deep neural network (DNN) was used to build a classifier to discriminate CAG expanded, *HTT* KO from controls on images from fixed and stained cortical organoids containing clusters of cells^30^. We wanted to harness the power of ML, but at a single cell resolution and over time. Therefore, we incorporated an innovative way to capture dynamic changes resulting from the CAG repeat expansion or loss of HTT that would otherwise be missed at a single timepoint. To do this, we subjected images of live eCNs to feature extraction and object recognition algorithms ^31,32,33,34,35,36,37,38,35^ that capture a range of engineered features commonly used in shallow machine learning approaches^39,40^. We analyzed how these features evolved over time as a proxy for monitoring dynamic cellular changes during development. Our results highlight those that are distinct versus overlapping morphologic alterations in response to the CAG expansion compared to the *HTT* KO.

Pluripotent stem cells (PSCs,patient derived and isogenic) have been used to model transcriptional and epigenetic dysregulation caused by CAG repeat expansion in various neural and neuronal populations. Dysregulation was observed in pathways associated with axonal guidance^26,41^, TGFβ, WNT, and BDNF signaling^41–43^ and neurogenesis particularly via NEUR0D1^26,42^. HTT’s known role in many of these pathways including BDNF signaling^19^ and neurogenesis^20,44^ suggest that HTT loss of function is contributing to these differences. However, direct molecular comparison of dysregulation caused by CAG repeat expansion and HTT loss of function has yet to be examined. Here we use multi-omics analysis to model pathways of dysregulation in CAG repeat expansion and HTT KO and examine where they intersect or differ.

Further, we integrated this multi-omics data with our morphological analysis and discovered specific phenotypes contributed by *HTT* KO and CAG repeat expansion. Both *HTT* KO and expansion caused neurodevelopmental changes, however these comprise features and pathways that are both overlapping and distinct. By identifying key molecular drivers of these phenotypes and separating out pathways that are unique to KO versus CAG expansion, our data have important implications for future therapeutic development for HD.

## Results

### Cortical neuron differentiation and cell-type characterization across *HTT* CAG repeat and KO RUES2 lines

To better understand the cellular consequences of the *HTT* CAG repeat expansion versus loss of HTT function, we employed a systems approach using multi-acquisition omics analysis including ATAC-seq, ChIP-seq, RNA-seq and proteomics in conjunction with cellular phenotyping analyses (**Figure 1A-C**). To eliminate the patient-to-patient and cell line-to-cell line variability which complicates differential analysis, we applied these analyses to a previously reported isogenic series developed in the RUES2 ESC line^20^. The series use consists of control clonal lines with normal CAG repeat lengths (20CAGn1, 20CAGn2, 20CAGn3, and 20CAGn4), an expanded line harboring 56 CAG repeats (56CAG), three clones of a juvenile-onset CAG expansion containing 72 CAG repeats (72CAGn1, 72CAGn2, 72CAGn3), and one line with homozygous deletion of *HTT* exon 1 resulting in loss of the HTT protein expression(*HTT* KO)^20^ (**Figure 1A for repeats and Supplemental Table 1 for clone designations**). These lines were differentiated down a cortical neuron lineage to generate eCNs using a recently developed protocol based in part on Shi et al. 2012^45^ with the addition of a Sonic hedgehog (SHH) inhibitor, cyclopamine^46^, to direct away from a medial ganglionic patterning^47^, and extended FGF2 signaling to enhance the derivation of glutamatergic cortical neurons. Maturation was accelerated by using the gamma secretase inhibitor DAPT, which forces cells to exit the cell cycle^48^ (**Figure 1B**). After ∼ 35 days of differentiation, these eCN populations consist of a large portion of MAP-2 positive cells as well as those expressing the forebrain marker FOXG1, early cortical lineage marker (layer VI) TBR1, and a small number of cells positive for cortical layer V marker BCL11B (**Figure 1D and Supplemental Figure 1).**

**Figure 1.**
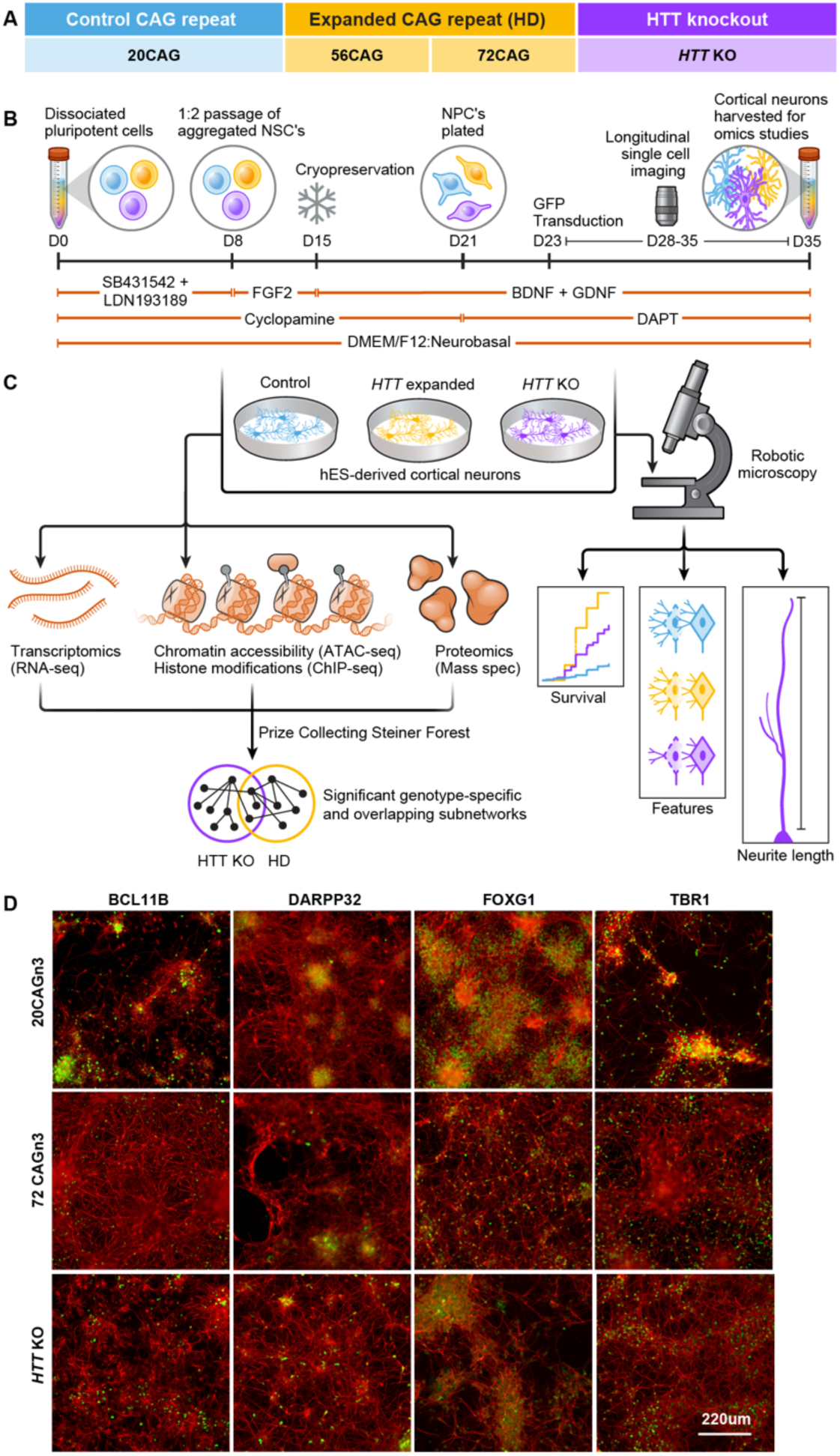
Methodological overview and study design. (A) Table of isogenic cell lines used in this study, *denotes the parental line modified to generate all others. A complete list of cell lines can be found in **Supplemental Table 1** (B) Schematic overview of the cortical neuron differentiation depicting the timeline of study measurements. The top bar denotes the differentiation timeline, bottom bars delineate the small patterning molecules used during each period of the protocol. (C) Schematic overview showing differentiation into cortical neurons (eCNs) used for robotic imaging (RM) analysis as well as generation of the 4 different omics assays (RNAseq, ATACseq, ChIPseq, and Mass Spectrometry), and integration of all data sets to reveal dysregulated pathways using Omics Integrator. D) Representative immunofluorescence images showing expression of neuronal markers in the eCNs that were subjected to RM and fixed at differentiation day ∼35. Expression of BCL11B, DARPP32, FOXG1, Ki67 and TBR1 in HTT KO, 20CAG and 72CAG eCNs. CTIP2, DARPP32, FOXG1 and TBR1 (green) were co-stained with MAP2 (red) and Ki67 (green) was colocalized with nuclei (blue). scale bar = 220◻m

We examined the differentiation efficiency of each line by staining for cell-specific markers at differentiation day ∼35. eCNs were fixed, stained and quantified to ascertain the percentage of positively labeled cells for each antibody using a modified Cell Profiler pipeline as previously described^49^. All clones (20CAGn3/n4, 72CAGn1/n2/n3 and *HTT* KO) of eCNs patterned to contain ∼60% FOXG1 and ∼30% TBR1 positively expressing cells (**Supplemental Figure 2A-D).** We did not detect a significant difference in the numbers of these cell types across all clones. These cultures developed a small proportion (∼10%) of inhibitory GABAergic neurons as identified by neurons expressing DARPP-32. There was a subtle, but significant reduction of the number of DARPP-32 positive cells in the *HTT* KO compared to the 20CAG eCNs (**Supplemental Figure 2E, F).** We also observed a small proportion (∼10%) of BCL11B positive cells, with the *HTT* KO displaying slightly lower number of positively labeled cells compared to control eCNs (**Supplemental Figure 2G, H)** and a small portion of persistent proliferating cells, as identified by the Ki-67 marker (**Supplemental Figure 2I, J)**.

### Expanded repeat and KO eCNs display differences in survival, neurite length and morphological differences over time compared to controls

We next asked if CAG-expanded and *HTT* KO eCNs displayed differences in cell health compared to control eCNs. Previous reports found that CAG-expanded neurons displayed a higher rate of death than controls.^25,50^. To assess survival, we transduced eCNs with the morphology marker EGFP driven by the Synapsin promoter (Synapsin:EGFP from Signa Gen) on differentiation day ∼24, and imaged them daily using robotic automated longitudinal imaging (Robotic Microcopy [RM]) (**Figure 1B,C, Supplemental Figure 3A**) to determine the cumulative risk of death of each line as previously described^25,50–60^. Fluorescent eCNs displayed typical neuronal morphology with an oval-shaped cell body and numerous axonal or dendritic processes (**Supplemental Figure 3A)**. We observed a large range of death rates across all lines and experiments (**Supplemental Figure 3B** therefore we used a Cox proportional hazards mixed effects model^61^ that accounts for batch, clone as well as unknown variability, to compare groups. This statistical approach showed that eCNs that contain 72 CAG repeats (72CAGn1/n2/n3) die significantly faster than control eCNs (20CAGn3/n4) (Hazard ratio (HR) = 2.2, p=3.7 E^-05^). There was no statistically significant difference between the controls and the *HTT* KO eCNs (**Figure 2A**).

**Figure 2.**
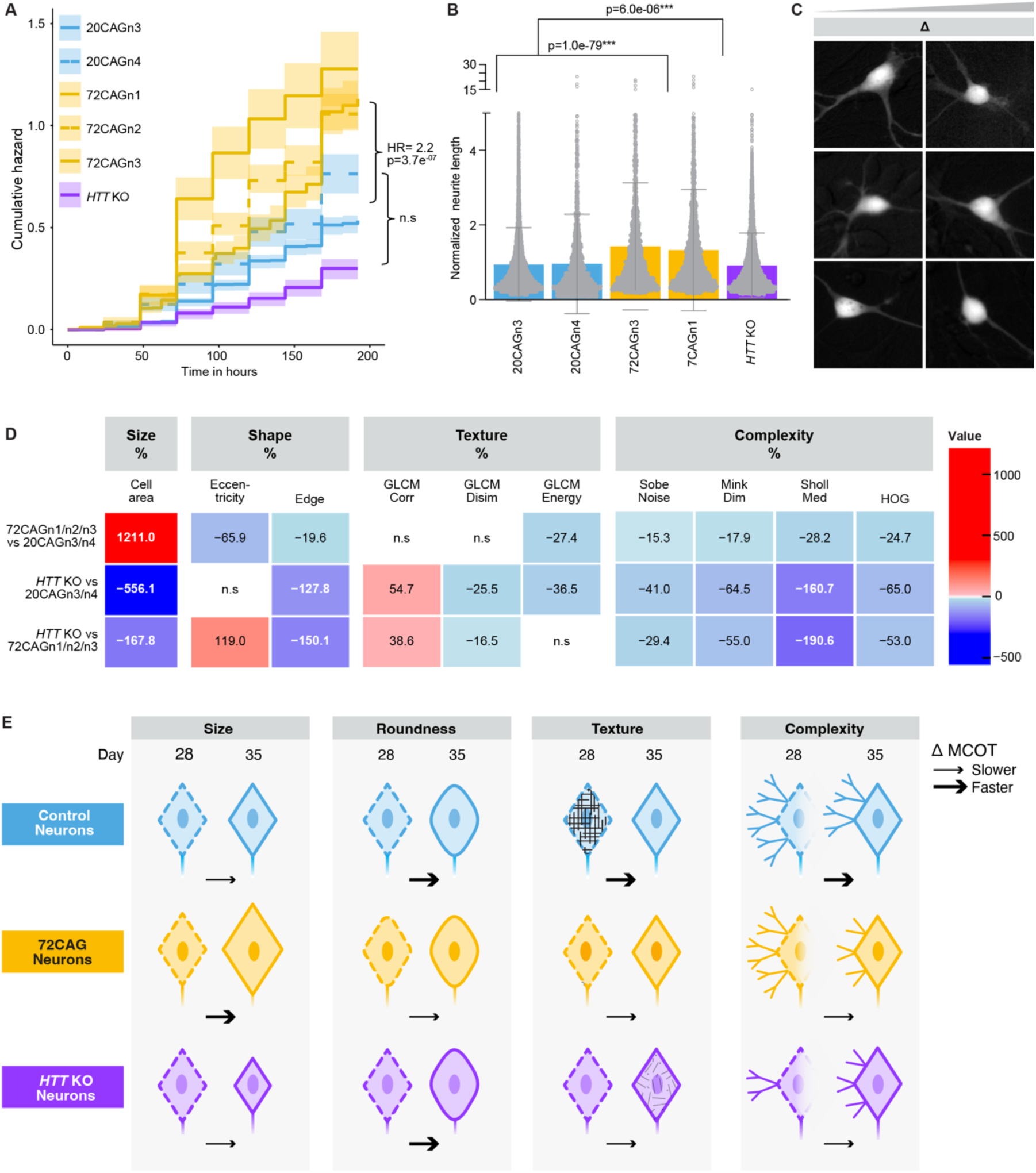
Live cell imaging of eCNs shows spontaneous degeneration and morphological changes between CAG-expanded HTT knockout and control eCNs. differences in neurite length across *HTT* genotypes. **A)** eCNs were transduced with LV-Synapsin-EGFP and tracked for 7 days as shown in example images in supplemental Figure 3. Using manual curation, cells were determined live or dead, and the cumulative death rate (live/dead cells) was plotted over time as previously described. A Cox mixed effects model^61^ was used to calculate hazard ratio of 72CAG eCNs cells to controls, HR= 2.12 with p-value of 3.7e-07 ***. There was no difference between control eCNs and KO eCNs, HR=1.2, p=0.43. 20CAGn3 = 1432 neurons; 20CAGn2= 515 neurons; 72CAGn1= 575 neurons, 72CAGn2 = 972 neurons; 72CAGn3= 340 neurons, *HTT* KO= 555 neurons, 9 experiments. **B**) Neurites were measured using CellProfiler to capture neurite length on images acquired by RM differentiation day ∼32. Neurite values were normalized as previously described^26^. We used a LMM^123^ to compare groups. Neurites from 72CAG eCNs are longer than control and KO eCNs, p= 1.0e^-79^ whereas neurites from the HTT KO are shorter than control eCNs p= 6.0e^-06^ 20CAGn3= 10,849 neurons, 20CAGn4= 2725 neruons, 72CAGn3 = 2320 neurons; 72CAGn1= 4092 neurons, *HTT* KO= 9,691 neurons, from 11 experiments. **C**) Representative images show changes in cortical neuronal morphology over time for the 20CAG, 72CAG and *HTT* KO genotypes. **D**) Time interaction feature analysis reveals dynamic morphological changes in CAG expanded and *HTT* knockout eCNs. Heatmap showing changes across the different groups over time. A time interaction model was used to determine the change in feature values across time from differentiation day 28–35 for each group. The MCOT is not a rate, but a coefficient that is described in detail in the methods section. Top, 72CAG vs 20CAG, middle KO vs 20CAG and bottom KO vs 72CAG. The values listed in the table denote the interaction between the time variable and the disease status variable scaled by the change over time in the reference (72 CAG or *HTT* KO versus control) and expressed as percentages. The color reflects the degree of change between the two groups and the value in the middle is the percent change and the direction. For example, the area feature in 72CAG eCNs increases by 1211% compared to control eCNs whereas the KO is changing by 556% but in the opposite direction. n.s. = not significant. The delta represents passage of time between the images on the left and the images on the right. **E**) Illustration summarizing how each genotype shows differences in morphology and their respective rates of change with respect to time.

We next examined a second cellular hallmark, neurite length, which is correlated with neurodegeneration^62,63^ and neurodevelopment^3,26,51–59,64,65^. We quantified the length of each process from each eCN soma, as previously reported^66^, in images acquired from RM on differentiation day ∼32. We found that CAG-expanded lines displayed subtly longer neurites than the controls (**Figure 2B**). While this result appears to contradict the increased risk of death in the 72CAG eCNs (as dying neurons display contracting or blebbing neurites), it is consistent with previous observations from neurons from HD patient lines^26^ and other ESC-based models of HD *in vitro*^67^, and from neurons in the brain of early-stage HD patients^68–70^. In contrast, the *HTT* KO eCNs displayed subtle but significantly shorter neurites than controls (**Figure 2B**). Thus, the differences in neurite length may represent neurodevelopmental alterations across the groups as opposed to neurodegenerative ones.

To capture neurodevelopmental changes, we devised a novel platform using feature extraction commonly used in ML to examine morphological changes at a single instance or across time during neuronal differentiation and maturation between the control, CAG-expanded neurons and HTT KO neurons. D24 eCNs transduced with synapsin::EGFP were imaged as described above using RM and processed in our image-processing pipeline Galaxy software^71^ (**Supplemental Figure 3C)** and cropped so that only the cell body and immediate processes were captured on a cell-by-cell basis (**Figure 2C, Supplemental Figure 3C**). These cropped images were then subjected to our custom-built ML feature-based platform that is a collection of algorithms that relay information about structure on a cell-by-cell basis (**Supplemental Figure 3D)**.

At a single time point (differentiation day 28), we detected subtle, but highly significant and reproducible differences across a small subset of the features. For instance, the perimeter of each cell as captured by the “Edge” feature captures the perimeter of each object and relates to both size or complexity of an object^37^. The ‘Edge” in the 72CAG eCNs, suggesting a more complex shape compared to the controls (**Supplemental Figure 4A).** This is consistent with the observation that the 72CAG eCNs have longer neurites than controls (**Figure 2B**). Likewise, the “Edge” of *HTT* KO eCNs was smaller than controls, also consistent with observation that they have shorter processes (**Figure 2B**). The “eccentricity” feature captures of the roundness of the cell (e.g. round versus elliptical)^72^ showed that the 72CAG eCNs were rounder than the controls at this time point (**Supplemental Figure 4B**). We also observed that the median number of processes emanating per soma captured by the “median Sholl” feature was significantly different in the *HTT* KO eCNs compared to control and 72CAG eCNs (**Supplemental Figure 4C**).

However, these differences across groups at day 28 were small. Previous studies have reported developmental alterations in HD patient derived i-neurons^47,50,73–76^ and we wondered if similar changes occurred in the eCNs. Therefore, we examined if these features changed over time as a way to capture morphological changes that could reflect developmental shifts. eCNs that expressed synapsin::EGFP were imaged using RM as described above at differentiation day ∼28 and again at day ∼35 (**Figure 2C**). Within each group of eCNs, all features displayed robust differences from the first time point to the last time point suggesting that the morphology of eCNs changes over time during differentiation **(Supplemental Figure 5**). To estimate this shift over time, we modeled the shift across the two time-points due to a change in disease status via an interaction term^77,78^ we call the “Morphology Change Over Time” or “MCOT”(the time:disease_status interaction terms were scaled by the change over time in the 72CAG and *HTT* KO or control and expressed as percentages (**Figure 2D**).We evaluated features of eCNs that harbored the 72CAG or *HTT* KO compared to control eCNs. We observed that more than 1/2 of the features (11 features out of 18 total) displayed significant MCOTs in the 72CAG eCNs compared to controls, ranging from subtle (∼15%) to 12-fold (1200%). Further, the *HTT* KO eCNs also displayed robust MCOTs of features compared to control eCNs (**Supplementary Figure 5**).

The most notable features capture changes in cell size, shape, complexity and texture and are summarized in a heatmap in **Figure 2D**. For example, the “cell area” which captures the size of the cell was found to change at a larger rate in the 72CAG eCNs compared to controls which were more static. The increase was ∼1200% greater in the 72CAG eCNS compared to controls (**Figure 2D and Supplemental Figure 5A)**. In an opposite fashion the “cell area” decreased in the *HTT*KO compared to the 72CAG and controls (**Figure 2D and Supplemental Figure 5A)**. The change in cell size may reflect rates of cell growth that capture developmental trajectories that arise as a result of either loss of HTT as opposed to those that are caused by the CAG repeat expansion.

The MCOT for “edge” was larger in the control compared to 72CAG and *HTT* KO eCNs (as observed by a −22% overall change) suggesting that the control eCNs are becoming more complex faster. The changes were more dysregulated in the *HTT* KO eCNs compared to the controls or 72CAG eCNs (by −128% and −150% respectively) (**Figure 2D and Supplemental Figure 5C)**. The “eccentricity” feature at day 28 suggested that the 72CAG eCNs were rounder than the control cells, however the MCOT for eccentricity was higher in 20CAG eCNs compared to the 72CAG eCNs (as observed by a −66% difference) indicating that the control eCNs became rounder faster than 72CAG eCNs during this stage of development (**Figure 2D and Supplemental Figure 5D**). The *HTT* KO MCOT for “eccentricity” was comparable to the control but was significantly different from the 72CAG eCNs, suggesting that this feature may capture a morphological change that is specific to the CAG expansion and may be interpreted as a “gain-of-function” property (**Figure 2D and Supplemental Figure 5D)**.

Other features capture information about an objects’ texture such as the Gray-Level Co-occurrence Matrix (GLCM)^79^ which measures the spatial relationships between neighboring pixel intensities. We observed that the “GLCM energy” feature ^80^, increased in all lines over in time in culture, suggesting that that the eCNs smoothen and become less textured as they differentiate. The MCOT of the 72CAG eCNs differed from controls by ∼ 27% showing less change over time (**Figure 2D and Supplemental Figure 5F**). There was no difference between the 72CAG compared to the *HTT* KO eCNs, suggesting that this feature may be capturing a cellular rate of change that reflects a loss of function due the CAG expansion. MCOTs for other GLCM-like features such as GLCM Dissimilarity (which measures contrast between pixels) and GLCM Corr (measures the linear relationship between pixel pairs)^80^, were only significantly different in the *HTT* KO eCNs (**Figure 2D and Supplemental Figures 5G,H**). This suggests that these features capture a true loss of HTT function that is distinctly different from the loss of function that occurs from the CAG repeat expansion.

Additional features capture information about an objects’ complexity such as “HOG” (histogram of gradients)^36^ or Minkowski-Bouligand Fractal Dimension (Mink Dim)^81^, which has been used to detect differences in complexity in human tissues^82^, and the “sobel noise” which is an edge detector based on gradient changes across an image^83^. The sholl intersection^84^ feature captures the number of processes protruding from a soma which can indicate the pattern of neurite outgrowth. The majority of “complexity” features all decreased over time all groups, suggesting a decrease in cell-shape complexity as the neurons matured. However, the MCOT for complexity features decreased more in the control eCNs compared to the 72CAG or *HTT* KO eCNs (**Figure 2D and Supplemental Figure 5 I, J, K, L)**. There are other features that display differences across the control, 72CAG and *HTT* KO eCNs (**Supplemental Figure 5B, E, M, N,O,P Q),** but the biological meaning of these features is unclear.

To summarize, 72CAG eCNs grow larger faster than control eCNs, which are more static, whereas *HTT* KO eCNs grow more slowly and stay smaller than both the 72CAG and control eCNs (**Figure 2D, E).** The texture and complexity of the neurons increases most robustly in the control eCNs compared to the 72CAG and *HTT* KO eCNs (**Figure 2D, E**). These results suggest that *HTT* CAG repeat length expansion and loss of HTT function both influence cortical neuron morphology and development, however the CAG repeat influences metrics in ways that cannot be explained by HTT loss of function alone.

### Overview of OMIC analysis of ESCs and eCNs

To explore the gene expression, protein and epigenetic profiles of the isogenic cell lines, we conducted transcriptomics (RNAseq), proteomics (mass spectrometry), and epigenomics (ATACseq and ChIPseq with H3K4me1, H3K4me3, H3K27ac and H3K27me3 histone marks) at both the pluripotent and eCN stage as described in Methods^85–88^. Two control cell lines (20CAGn1/n2), two expanded repeat lines (56CAG and 72CAGn1) and the *HTT* KO cell lines were collected across 3 successive passage replicates at the pluripotent stage to establish a baseline cell signature. Similarly, eCNs were generated in three sets of independent differentiations. Generated eCN’s were assessed to by ICC and in general, these cultures were relatively homogeneous with a higher percentage of MAP2-positive neurons and a lower percentage of NESTIN-positive progenitors and, on average, did not display large variability across cell line or experiment (**Supplemental Figure 6).** Cultures express forebrain lineage marker FOXG1 (**Supplemental Figure 6B) and** contain a high percentage of the mature neuronal marker MAP2 (**Supplemental Figure 6A)** (mean percent positive: 85.81, range: 67.5-99.16, and early cortical lineage maker TBR1 (mean percent positive: 48.90, range: 18.81-83.74) (**Supplemental Figure 6C**). eCN populations also expressed low levels of BCL11B (**Supplemental Figure 6D)** similar to those use for imaging studies. Pooled pellets from each line at the pluripotent and eCN stage were subjected to the same set of assays. Finally, we integrated the primary assay data using the OmicsIntegrator tool (**Figure 1C**), using the differential analyses as inputs to the Prize-Collecting Steiner Forest algorithm^85,89^. The algorithm identifies the highest-confidence protein-protein interactions involving differential omics from a reference of known protein-protein interactions.

### HTT KO has the strongest omics effect in pluripotent cells, and eCN stage cells display clustering based on CAG length and HTT KO

Each of the pluripotent omic data sets (RNAseq, ATACseq, ChIPseq, and DIA-mass spectrometry) were subjected to unbiased clustering analysis prior to differential analyses. Principal component analysis (PCA) for transcriptional data showed that all growth replicates of each cell line clustered closely and the 20CAGn1 and 20CAGn2 lines clustered together; however, the two CAG-expanded lines (56CAG and 72CAGn1) did not cluster together (**Supplemental Figure 7A**). The minimal separation of the 20CAG lines from the CAG-expanded lines likely reflects a minimal effect of CAG-expansion on cells at the pluripotent stage. The most highly expanded line, the 72CAGn1 line, had fewer differential transcriptomic changes and differential ATAC-seq peaks compared to the controls (20CAGn1 and 20CAGn2) relative to the number of differentially expressed genes and differentially accessible peaks between *HTT* KO cells and controls as well as the 56CAG cells and controls **(Supplemental Figure 7B, Supplemental Table 4, 5A-U**). Data integration analysis using the Omics Integrator tool showed one cluster of particular interest associated with *HTT* KO (**Supplemental Figure 7C**). In this cluster, the HTT protein is a node connected to its known interactor CREBBP/CBP^90^ as well as its downstream target, FOS. The HTT protein also interacted with TBP, which acts in a complex with CREBBP and HTT for transcription-coupled repair^91^. Additional nodes TOP2A and FOXM1 potentially highlight HTT’s normal role in DNA replication and repair.

Alterations in expression and epigenetics were next examined at the cortical neuron stage. We utilized PCA to explore the clustering patterns of gene expression of the differentiated eCNs using the OMIC data. Both CAG expansion and loss of HTT function displayed significant effects on gene expression, protein levels and epigenetic readouts in the eCN population. CAG repeat expansion and HTT loss accounted for the majority of variance by PCA, however some variance was observed between replicates, likely due to heterogeneity across independent differentiations. Control eCNs clustered across all data sets and all cell lines separated by genotype, particularly for the RNAseq, ATACseq, and H3K4me1 ChIPseq data sets (**Supplemental Figure 8A-D**). In the RNAseq data, CAG repeat seemed to account for the variability across PC1, with the 72CAG line clustering at one end and the control lines clustering at the other (**Figure 3A**). The first principal component accounted for 80% of the variance. For the ATAC-seq peaks and protein expression in eCN cell lines, we observed separation by the CAG expansion for both 56CAG and 72CAGn1 along PC1, while in the H3K4me1 peaks, we observed this separation along PC2 (**Supplemental Figure 8B-D**). In the PCAs, *HTT* KO cells tended to cluster with the CAG-expanded repeat cell lines, suggesting some common changes. We also performed PCA on other ChIP-seq data sets, however, the other epigenomic marks did not show separation by repeat length (**Supplemental Figure 8E-F**).

**Figure 3:**
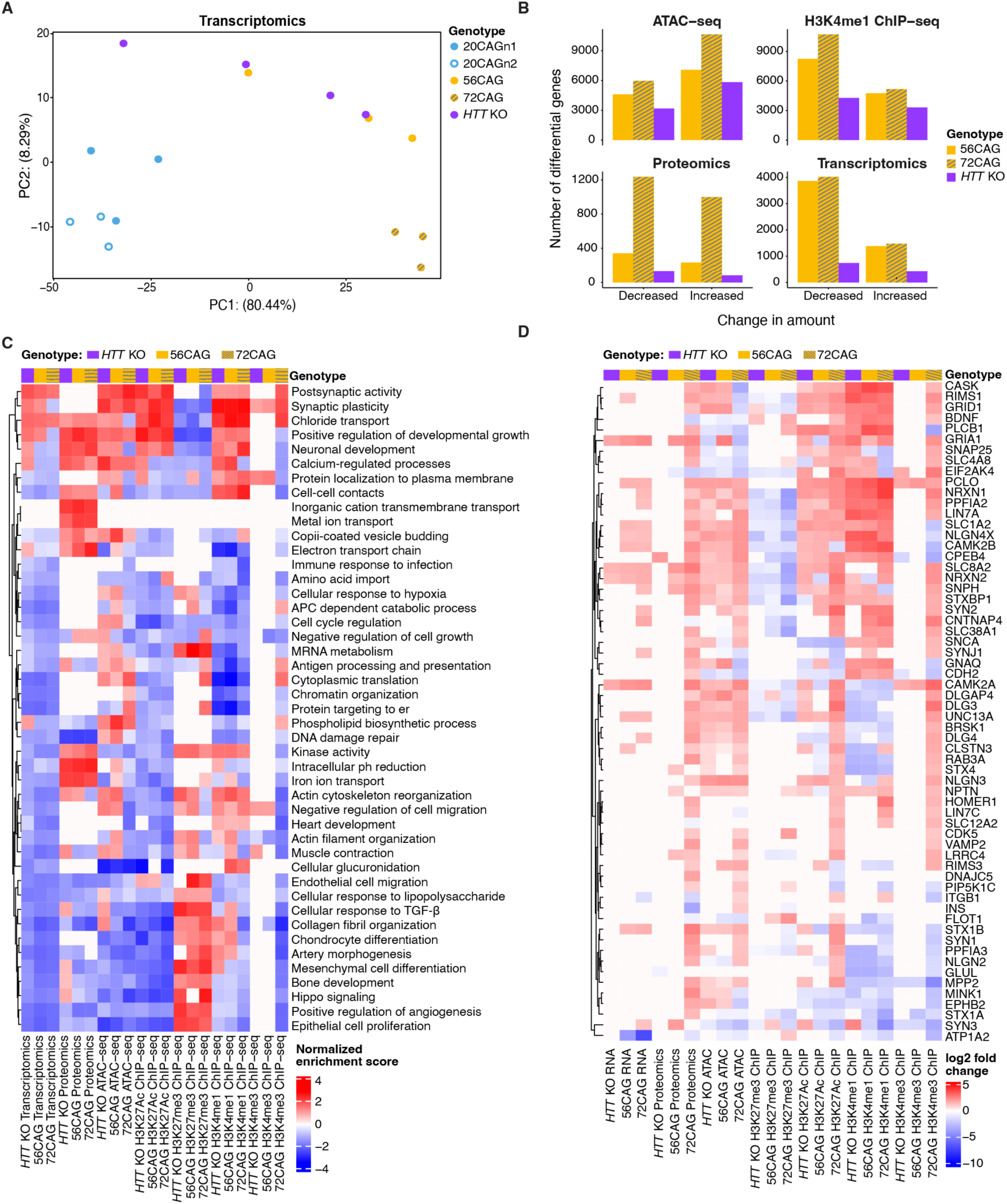
Analysis of primary assay data shows similarities across data types and between HD cells and *HTT* knockout cells. (A) PCA of cortical neuron transcriptomics shows partial separation by CAG expanded status along principal component 1. (B) Number of differential genes in ATAC-seq, H3K4me1 ChIP-seq, proteomic and transcriptomic data comparing 56CAG eCNs to control neurons (Gold bar), 72CAG eCNs (Gold bar), and *HTT* KO to control neurons (Purple bar) (C) Heatmap of the normalized enrichment scores of the union of significantly enriched pathways in transcriptomic, epigenomic and proteomic data in 52CAG eCNs, 72CAG eCNs, and *HTT* KO eCNs. Many pathways have the same direction of change in all three genotypes in each assay. Moreover, there exist pathways that are upregulated in all assays but downregulated in H3K27me3 ChIP-seq, which is a repressive epigenetic mark. (D) Heatmap of the log_2_ fold changes of the union of differentially expressed genes across assays and genotypes that are involved with postsynaptic activity. The genes that are displayed in this pathway were differentially expressed genes in at least one comparison of genotypes and assays. Many of these genes show the same direction in all omics except the repressive histone mark H3K27me3 ChIP-seq, which changes in the opposite direction.

### Pathway analysis and multi-omic differential analyses suggest coordinated changes across HTT genotypes

In eCNs, we observed 4,084 differentially expressed genes (DEGs) and 5,506 DEGs of the 56CAG and 72CAGn1 compared to controls, respectively (**Figure 3B**, **Supplemental Table 6, 7,** FDR<0.01, logFC>|1|). The number of DEGs for the *HTT* KO eCNs compared to controls were far fewer at 1168 (**Figure 3B, Supplemental Table 6, 7** FDR<0.01, logFC>|1**|**). Many of the DEGs in CAG-expanded eCNs overlapped with those identified in previous studies comparing HD to control iPSC-derived cortical^42^ and striatal neuron^50^ populations (**Supplemental Figure 9A,B**, hypergeometric p-value=2.0*10^-6^ and hypergeometric p-value<10^-16^ respectively). There was also significant overlap between the DEGs we identified in *HTT* KO eCNs compared to published results in iPSC-derived striatal neurons^50^ (**Supplemental Figure 5C**, hypergeometric p=3.8*10^-4^) and partial overlap with published results in cortical neurons (**Supplemental Figure 9D**, hypergeometric p=0.02). The overlap between DEGs identified in RUES2 lines and those found in iPSC-derived neurons from HD patients supports the utility of the isogenic eCN-HD model and its ability to discriminate between loss-of-function versus gain-of-function cellular changes attributable to the CAG expansion.

Similar to the transcriptomic data, we identified 573 and 2,234 differentially expressed proteins (p<0.05, logFC>|0.6|) in the 56CAG and 72CAG repeat lines compared to control eCNs, but only 217 differentially expressed proteins comparing *HTT* KO to control eCNs (**Figure 3B**). The ATAC-seq and ChIP-seq data also had a greater number of differential peaks for the CAG expansion comparisons (**Figure 3B, Supplemental Table 6, 7**). Specifically, we found 11,676 and 16,587 differential ATAC-seq peaks for the 56CAG and 72CAGn1 repeat lines compared to controls (**Figure 3B, Supplemental Table 6, 7**).

Pathway analysis for each of these comparisons was performed across all data sets by gene set enrichment analysis (GSEA). Not surprisingly, considering HTT’s putative role in neurodevelopment^15,92,93^ and the early stage of the cortical neurons analyzed, several pathways relating to development and proliferation were enriched across multiple genotypes and assays (**Figure 3C**). These included Hippo signaling, developmental growth, and cell cycle regulation. For example, developmental growth regulation was enriched in *HTT* KO, 56CAG and 72CAG proteomics, *HTT* KO, 56CAG and 72CAG H3K27me3 ChIP-seq, and *HTT* KO and 72CAG H3K27Ac ChIP-seq. Other pathways of interest included DNA damage repair, synaptic plasticity, chromatin organization, and negative regulation of cell migration (**Figure 3C**). The implicated DNA damage repair genes included important enzymes such as PARP1, XRCC1 and OGG1 as well as replicative enzymes like PCNA, POLE and POLD1, the latter identified as a candidate modifier of age-of-onset in HD^94^.

Some pathways had related enrichment scores across *HTT* genotypes. For example, some pathways that were significantly upregulated in *HTT* KO eCN transcriptomic data were also upregulated in the CAG-expanded eCNs by transcriptomic analysis (**Figure 3C**). Some of these enrichments in the transcriptional data were mirrored by significant enrichments in proteomics, ATAC-seq or activating ChIP-seq marks, and significant down-regulation of the repressive H3K27me3 mark (**Figure 3C**). A significant proportion of genes that were differentially altered in each assay had the same direction of fold change in *HTT* KO and CAG-expanded eCNs, designated as “concordant changes” (**Supplemental Figure 10A** FDR-adjusted hypergeometric test p-value less than 0.1). As an example, genes involved in postsynaptic activity showed coordinated changes across multiple genotypes (**Figure 3D**). Several genes had concordant changes across genotypes in more than one assay (**Supplemental Figure 10**). The presence of these concordant changes across *HTT* genotypes and assays supports the hypothesis that some of the pathways are similarly regulated in *HTT* KO and CAG-expanded repeat neurons, and that CAG repeat length expansion may at least in part cause reduced normal function of HTT.

### Network analysis shows similarities between biological processes in CAG-expanded and HTT KO eCNs

We used the differential signals between CAG-expanded eCNs vs. control eCNs to identify disease-relevant subnetworks. Specifically, we used the results of differential analyses comparing 56CAG to control and 72CAG eCNs to control as inputs to OMICS integrator^89^ which, as described above, identifies the highest-confidence protein-protein interactions from a reference of known protein-protein interactions. We refer to nodes that were not identified in our set of differential omics but were included in the network solution as “predicted nodes”. Louvain clustering of our pruned protein-protein interaction network showed subnetworks comprised of proteins that participate in core cell signaling pathways such as cell fate commitment, Hippo signaling, response to oxidative stress, histone methylation, extracellular matrix, cell cycling, and WNT signaling among others (**Figure 4A**). Many of these processes have been previously associated with HD. Support for these enriched pathways come from multiple lines of evidence across omics modalities (**Figures 4B-D**). Examples of subnetworks enriched for HD-associated processes that are supported by interactions among enriched proteins, epigenetic motifs and differentially expressed genes include those enriched for response to reactive oxygen species, nitrogen cycle metabolic processes and DNA-initiated DNA replication. In these subnetworks, at least half of the differential signals were seen in comparisons between 56CAG cortical neurons vs. control as well as 72CAG cortical neurons vs. control (**Figure 4B-D**). These results suggest that increased *HTT* repeat length is associated with reduced levels and potentially reduced activity of proteins involved in biological processes previously associated with HD (**Figure 4**).

**Figure 4:**
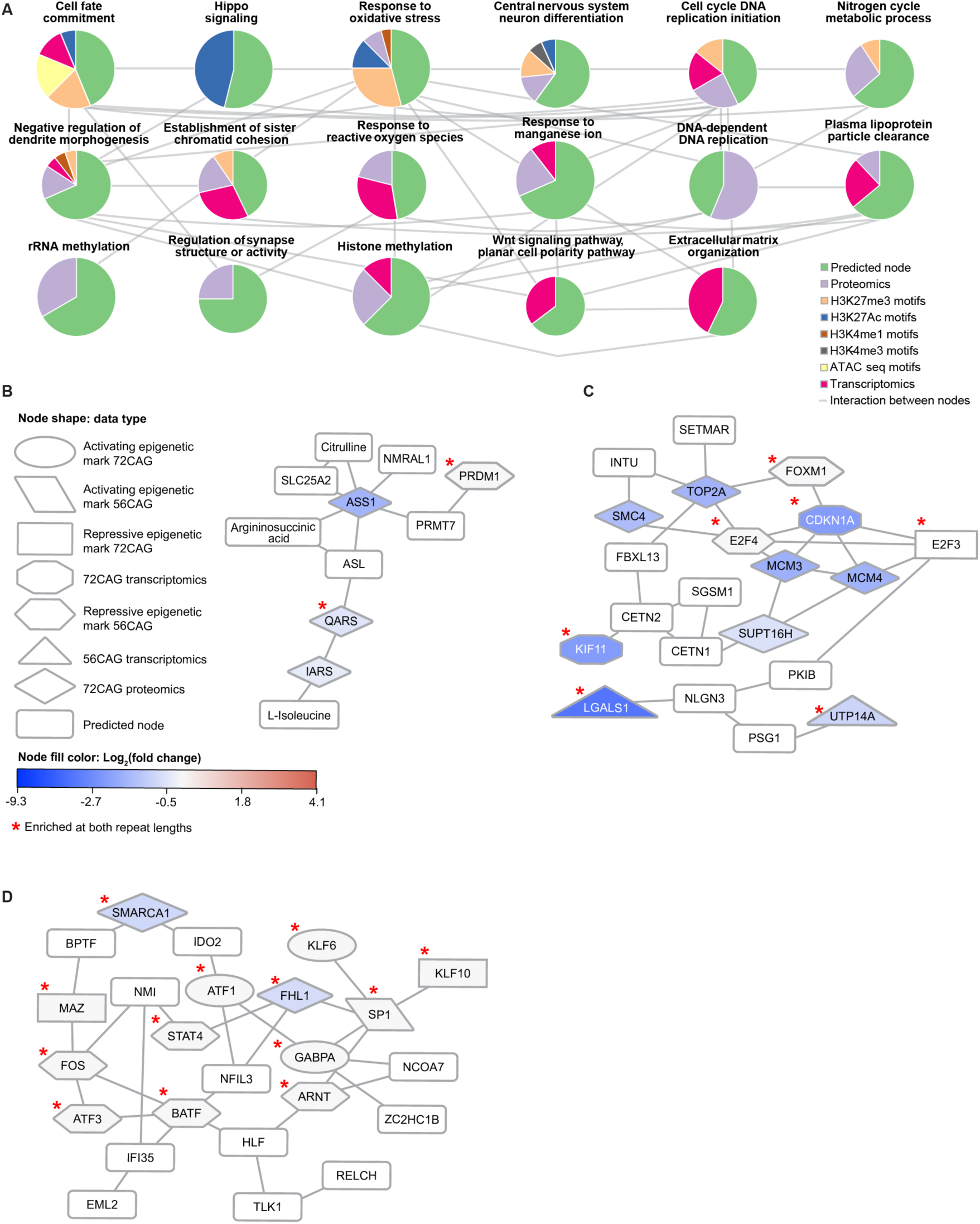
Mutli-omic network integration of CAG expansion-associated-Omics. (A) Multi-omic integration results for proteomics, epigenetics and transcriptomics enriched in 56CAG and 72CAG cortical neurons compared to controls. Each pie chart represents the origin of the nodes in the subclusters and is labeled with the enriched GO Biological Process. Edges between the pie charts indicate whether at least one node in one subnetwork interacts with at least one other node in the other subnetwork. (B) Subnetwork enriched for nitrogen cycle metabolic processes. Red asterisks indicate nodes that are enriched in the 72CAG and 56CAG cells. (C) Subnetwork enriched for DNA-dependent DNA replication processes includes nodes that are necessary for the DNA damage response process. Red asterisks indicate nodes that are enriched in 72CAG and 56CAG cells.

We also performed network-based multi-omic integration for data from *HTT* KO eCNs, revealing that some subnetworks were enriched for processes relevant to neuronal development, including axon guidance, NOTCH signaling, cell fate commitment and regulation of cell growth (**Figure 5A**).

**Figure 5:**
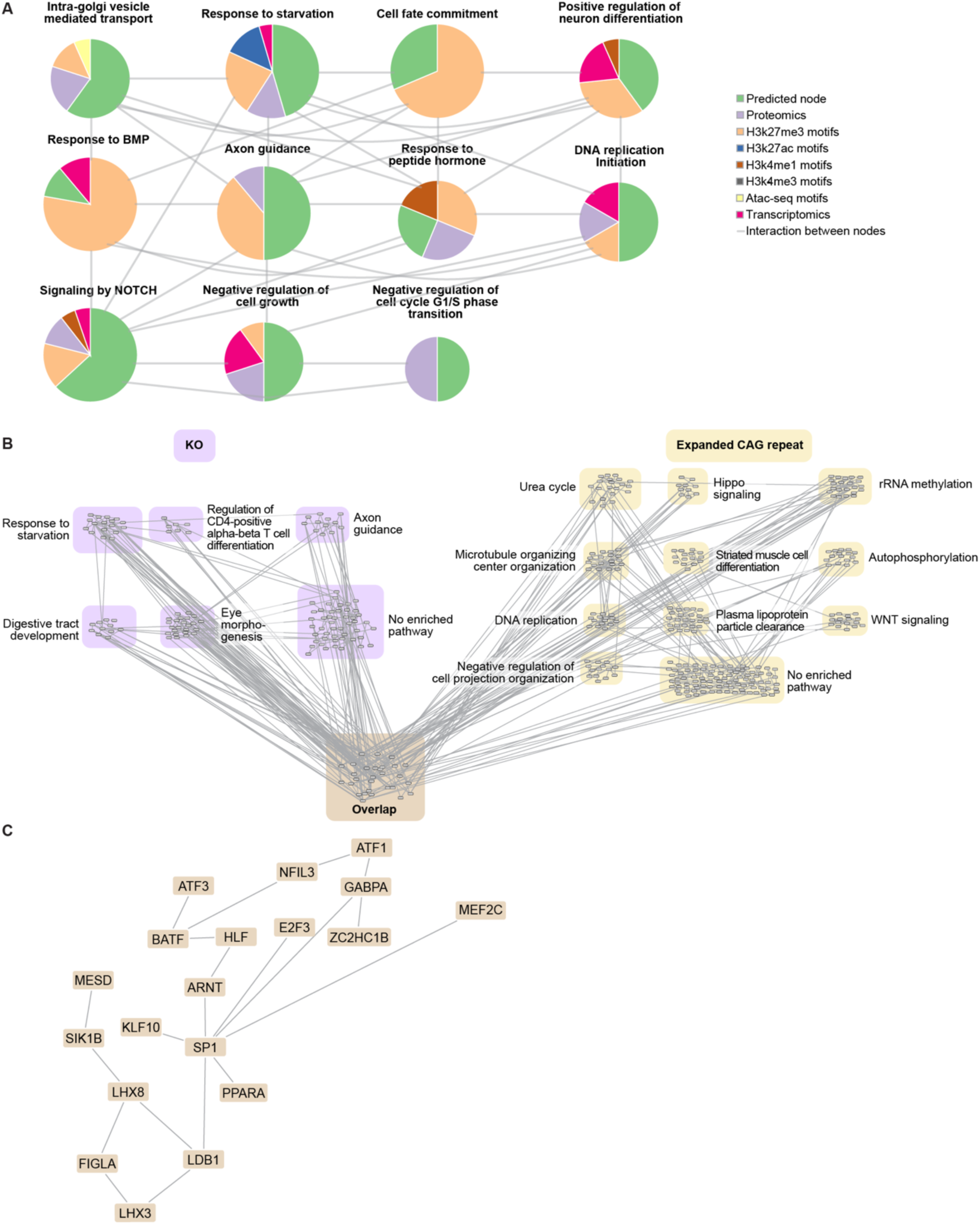
Integrative network analysis of *HTT* KO multi-omics show distinct and shared nodes and biological processes to the CAG expansion network. **(A)** Louvain clusters of the integrative network of differential proteomics, epigenomics and transcriptomics between *HTT* KO and control. The subnetworks are indicated by a pie chart and are labeled by the enriched GO Biological Process. Pie charts indicate the proportion of nodes represented by a given molecular data type. Edges between the pie charts indicate whether at least one node in one subnetwork interacts with at least one other node in the other subnetwork. **(B)** The union of nodes in the *HTT* KO and expanded CAG networks show that there exists a significant overlap in the nodes in the *HTT* KO and expanded CAG networks. The union of nodes between the two networks were re-clustered and labeled with enriched biological processes, with the KO nodes highlighted in purple, the HD nodes highlighted in gold, and the overlapping nodes highlighted in orange. The overlapped nodes are enriched for targets of MAPK signaling. **(C)** The largest connected component of the nodes and edges that are at the intersection of the *HTT* KO and HD networks.

Some of the processes enriched in the *HTT* KO network were also enriched in the CAG-expanded network, including neuronal differentiation, cell fate commitment and DNA replication initiation. To understand the degree of overlap, we visualized the union of the two networks (**Figure 5B**), clustering the unique and overlapping components separately. There were 29 nodes shared between the *HTT* KO and CAG-expanded repeat networks, which is greater than the number expected by chance between two networks of similar size (**Figure 5C, Supplemental Figure 11A, Supplemental Table 8)**. The overlapped network also had a larger connected component than any intersection of two random networks of the same size, supporting the hypothesis that the intersection of these two networks represent biological pathways as opposed to disconnected but shared nodes (**Supplemental Figure 11B**). Among the nodes that overlapped are MEF2C, ATF1, and MEF2A, which are targets of MAPK signaling^95,96^. Neuronal development-associated nodes were also found in this intersection such as SP1 and LHX3. The overlapped network highlights possible biological processes that are affected due to a loss of HTT function that occurs either as a result of a total loss of the protein or as a result of the CAG expansion.

The networks also revealed pathways specific to each genotype. *HTT* KO-specific nodes were enriched for starvation response and axon guidance, while the CAG-expanded exclusive nodes were enriched for Hippo signaling and WNT signaling as well as metabolic processes such as the urea cycle and plasma lipoprotein particle clearance (**Figure 5B**). These distinct networks may reveal pathways that are unique to the gain of functional activity stemming from the expanded CAG repeat versus.

Interestingly, in some cases, the CAG-expanded and *HTT* KO networks reflected similar processes, even when there were few overlapping nodes. Both networks showed an enriched H3K27me3 motif for the DNA-binding protein LHX3. We found that the LHX3 protein interacted with other nodes involved in neuronal development (**Figures 6A, 6B**). LHX3 is a homeodomain protein essential for nervous system development including of motor neurons, interneurons and pituitary development^97^. In the CAG-expanded networks, LHX3 interacted with the predicted nodes containing LHX6 and LDB1 that are involved in neuronal development (**Figure 6A**). Similarly, in the *HTT* KO network, LHX3 interacted with LHX4, LDB1 and ISL1, which are involved in neuronal differentiation^98^ (**Figure 6B**). Like LHX3, the ISL1 motif was enriched in the H3K27me3 ChIP-seq data. H3K27me3 is a repressive epigenetic mark, suggesting that both LHX3 and ISL1 are associated with reduced gene expression activity in *HTT* KO (**Figure 6B**). In the KO network, LHX3 was in the same subnetwork as CWF19L1, which is a gene commonly associated with spinocerebellar ataxia (**Figure 6B**).

**Figure 6:**
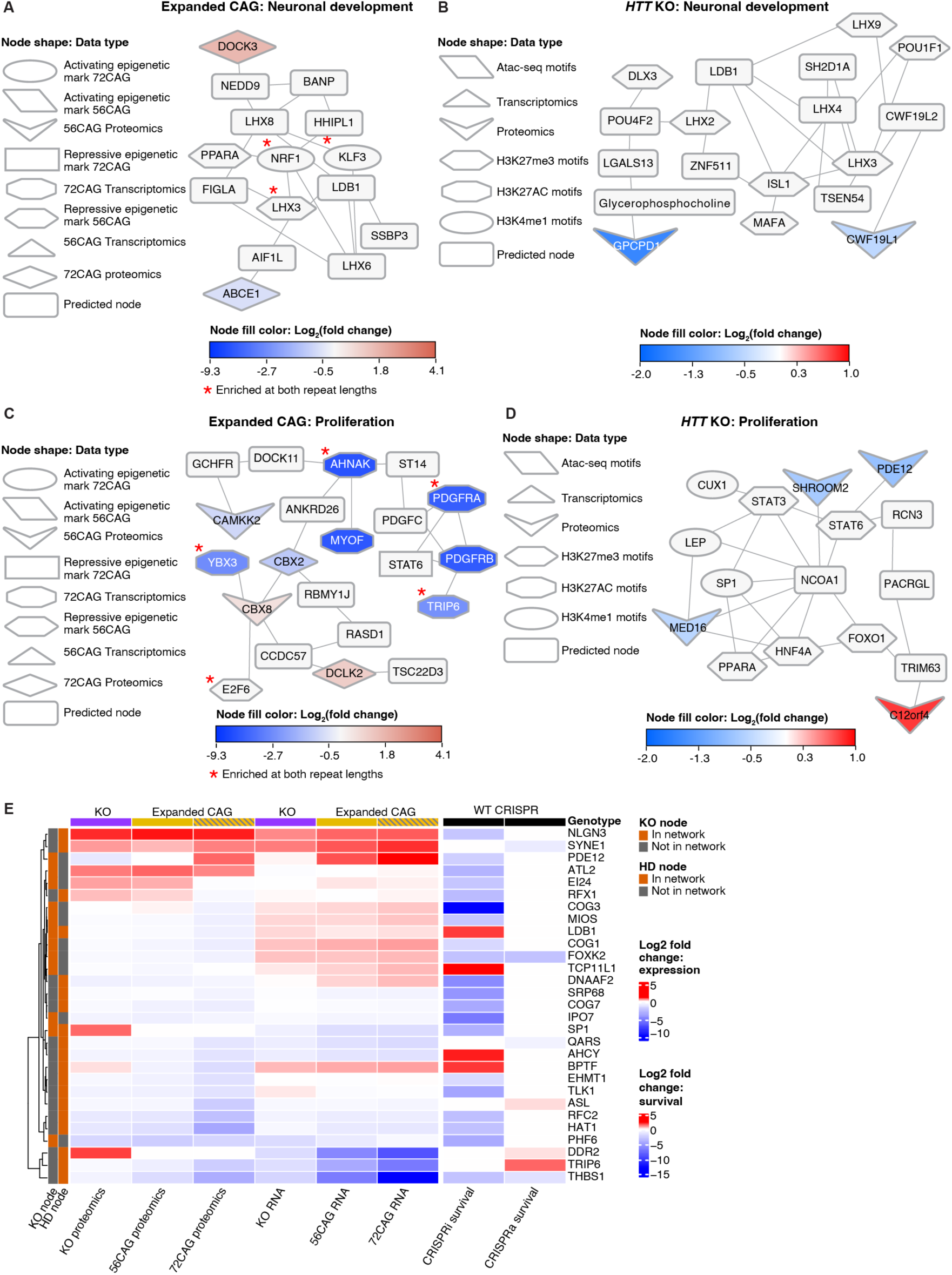
Subnetworks in the expanded CAG repeat and *HTT* KO networks show interactions that implicate similar biological processes despite differing node identities. (A) an expanded CAG subnetwork shows interactions between LHX3, LHX6, and LDB1, which are nodes associated with neuronal development. (B) An *HTT* KO subnetwork shows interactions between LHX3, ISL1, LHX4, and LDB1, which are nodes associated with neuronal development. (C) A expanded CAG subnetwork shows interactions between CBX8 and E2F6, which are nodes associated with cellular proliferation. (D) In an *HTT*KO subnetwork, we observe associations between MED16, HNF4A and FOXO1. HNF4A and FOXO1 are nodes associated with proliferation. Red asterisks in (A) and (B) indicate nodes that are enriched in 72CAG and 56CAG eCNs. (E) Heatmap showing the changes in the multi-omic data for nodes that appear in the *HTT*KO network or the CAG expanded network and significantly influence glutamatergic neuronal viability in a CRISPRi or CRISPRa viability in previously published data^101^.

We also observed a subnetwork in the CAG-expanded network that showed interaction between CBX8, a transcriptional repressor, and E2F6, a proliferation-associated transcription factor (**Figure 6C**). Since CBX8 is upregulated in the CAG-expanded eCNS as seen by proteomics and E2F6 is a motif associated with the repressive H3K27me3 mark, this subnetwork suggests associations with reduced proliferation and increased CBX8-mediated transcriptional repression. Similarly, we observed associations between MED16 and proliferation-associated transcription factors such as HNF4A and FOXO1 in the *HTT* KO network (**Figure 6D**). MED16 part of the mediator complex, which coactivator of RNA polymerase II and promotes transcriptional activation^99,100^, and the proliferation-associated epigenetic motifs were derived from H3K27me3 ChIP-seq data (**Figure 6D**). MED16 was downregulated in the *HTT* KO proteomics data (**Figure 6D**). These results suggest that, despite involving different nodes, the *HTT* KO network and CAG-expanded network represent protein-protein interactions involving similar changes in the regulation of transcription and proliferation (**Figure 6C, D**).

To understand how the nodes identified in our network analysis might affect neurodegeneration, we next analyzed our omics data and previously published perturb-seq data that measured single-cell differential transcriptomic changes in iPSC-derived glutamatergic neurons after CRISPRi knockdown or CRISPRa activation of target genes^101^ (**Figure 6E**). Specifically, after identifying genes common between the perturb-seq study and our own network analysis, we assessed whether knockdown or activation of these genes was associated with altered neuronal survival. Indeed, genes that were down-regulated in CAG-expanded and in *HTT* KO eCNs, such as *RFC2*, *THBS1*, *HAT1*, *PHF6*, *IPO7*, and *COG7,* were previously found to be associated with reduced neuronal survival upon knockdown, suggesting that these genes may have a role in HD-associated neurodegeneration (**Figure 6E**).

Our analysis identified a few possible, if challenging, targets for therapeutic intervention. Knockdown of *BPTF*, *AHCY*, *TCP11L1*, or *LDB1* improves glutamatergic neuronal survival (**Figure 6E**). *BPTF*, *TCP11L1* and *LDB1* were also upregulated in the CAG-expanded and *HTT* KO transcriptomics (**Figure 6E**). However, at the proteomic level, BPTF, TCP11L1 LDB1 were downregulated except for BPTF in *HTT* KO, which may reflect changes to protein homeostasis or compensatory effects (**Figure 6E**). Differences in transcriptomics and proteomics can be a consequence of compensatory effects and feedback loops, or differences in degradation pathways among other reasons. It will be of interest to determine the effects of genetic perturbation of *BPTF*, *TCP11L1* and *LDB1* on HD and assess whether these interventions would be useful therapeutic approaches, thus reconciling differences in transcriptomic and proteomic data. Additionally, the integration of RM with omics analysis, could help provide more functional information by examining how perturbations affect noted differences in cellular survival, size, and eccentricity as well as changes in cell size and cellular complexity over time.

## Discussion

In this study, we used a systems approach including live cell imaging and multi-omics to interrogate epigenetic, transcriptomic, and proteomic changes associated with CAG expansion and loss of HTT using an ESC-derived isogenic series of *HTT* modified lines patterned into cortical forebrain-like neurons (**Figure 1)**. CAG-expanded and *HTT* KO eCNs differentiated into cultures containing high levels of TBR1 and FOXG1-expressing neurons with a small subset of inhibitory neurons (**Figure 1, Supplementary Figures 1,2)**. Using RM, we show that CAG-expanded eCNs displayed a neurodegenerative phenotype of dying faster over time compared to control and KO *HTT* eCNs. Our multi-omic integration analysis identified overlapping changes across genotypes as well as subnetworks unique to each genotype, highlighting changes in developmental pathways and others associated with CAG expansion and loss of HTT function. Furthermore, the robust gene expression changes across the groups manifest as morphological changes such as alterations in neurite length as well as robust dynamic cell-structure changes across time. We found subtle, but significant differences with longer neurite length in CAG-expanded eCNs, consistent with previous reports, while finding that neurite length in *HTT* KO eCNs was reduced in a distinctly opposite fashion (**Figure 2**). Our results uncover key molecular targets and pathways for subsequent perturbation studies in additional patient cell lines and other models and sets up a system to examine the effects of perturbations at the molecular and cellular level.

Integration of epigenomics, proteomics and transcriptomics data from iPSC-derived models of human disease has emerged as a powerful approach for mechanistic studies of complex diseases^85,102–105^. This approach is especially important for neurodegenerative diseases that display a cascade of aberrant expression of many genes and proteins, and dysregulation of multiple biological processes/pathways. We used a network-based analytical framework to understand the biological processes underlying *HTT* CAG repeat length expansion and HTT knockout in an isogenic series to avoid patient-to-patient variability. By comparing the genotype-associated networks, we could infer which molecular changes were unique to CAG expansion and which had the same effect as HTT knockout, which could be especially useful for testing targets for therapeutic intervention given that current interventions are focused on total or allele-specific HTT lowering. The CAG-expanded and HTT knockout network shared significantly more nodes than expected by chance, and had subnetworks enriched for similar biological processes such as cellular proliferation and neuronal differentiation (**Figures 4**, **5**, **6**). Morphological analysis also showed changes in the rates of cortical neuron development in CAG-expanded and HTT knockout neurons (**Figure 7**). We note that the common pathway enrichments between the CAG-expanded and HTT knockout networks were comprised of different nodes (**Figure 5**, **6**). The overlap between the two networks included MEF2C, ATF1 and MEF2A, which are targets of MAPK signaling (**Figure 5B**). Neither MAPK signaling nor the set of MAPK targets were enriched in the single-omic analyses, highlighting the power of the network analysis to identify new nodes and biological processes that represent shared processes between expanded CAG repeats and HTT KO.

**Figure 7.**
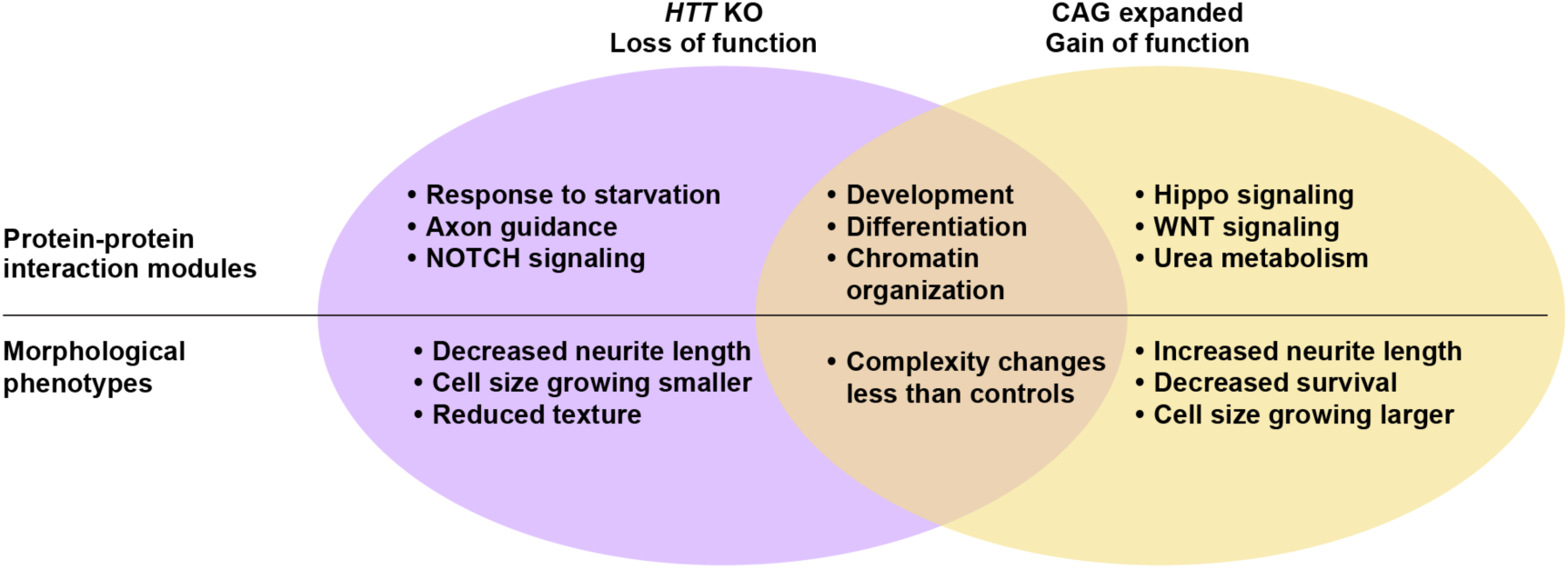
Schematic Illustration summarizing overlap and unique features of CAG repeat expansion and KO.

Early HD studies supported the idea that the CAG expansion led to gain-of-function characteristics that were toxic to cells. This included human genetic studies that observed that individuals with *HTT* deletion syndromes did not phenocopy the symptoms associated with typical HD (e.g.^18^). This was also supported by early mouse models where homozygous knockout of mouse huntingtin (Hdh) was embryonic lethal^106,107^, whereas heterozygous knockout mice survived to adulthood and were phenotypically normal^106^. Further, mutant HTT delivery is sufficient to rescue the KO mouse phenotype^108^. However, subsequent mouse models using conditional KO as well as cell culture models have led to a reevaluation of how reduced HTT function and timing of that reduction may contribute to disease progression^15^. Brain-specific inactivation of Hdh in mice results in motor deficits and striatal degeneration, similar to that seen in mouse models of HD^92^, as well as cellular hallmarks of impaired autophagy ^10^. iPSC and ES-derived cell models expressing expanded CAG have impairments in processes associated with normal HTT function such as impaired mitotic spindle orientation and transcription of BDNF^20,41,44,109^. Additionally, conditional silencing of Hdh in the mouse cortex affects excitatory synapse formation resulting in an excess of hyperactive excitatory synapses in the cortex early in development (p21) that later deteriorated by 5 weeks^93^. This same phenotype was found in the zQ175 mouse model of HD^93^. Interestingly, synaptic function was significantly dysregulated in our data sets. with several genes such as *GRIA1*, *CASK*, and *SCNA*, being upregulated in either the transcriptome, proteome, or epigenome in both CAG-expanded and HTT KO lines, suggesting that our data set likely reflects this early timepoint of dysregulation that leads to later progression of disease (**Figure 3**).

Furthermore, our network-based analysis identified developmentally relevant transcriptional factors SP1 and LHX3 in the overlapped subnetwork, further emphasizing neuronal development as an important process associated with HTT knockout and *HTT* CAG repeat length expansion. This observation is further supported by our analysis that there are dynamic morphological changes in eCNs (**Figure 2**). HTT is critical for neurodevelopment, and many deficits seen with HTT loss are recapitulated with CAG expansion, such as mitotic spindle dysfunction affecting cell fate particularly in cortical progenitors^44,110^ as well as a loss in the ability to self-organize into early developmental structures such as rosettes^20^. SP1 as a node in this shared network is not surprising, as several studies have evaluated not only its interaction with HTT ^111^ but also its potential role in HD progression^111–113^. Recent integrated transcriptional analysis of multiple neuronal lineage differentiations even proposed SP1 as a common regulator or dysfunction in HD neurodevelopment. Our findings here further support that there may be a role for SP1 in early dysfunction in HD.

Our expanded CAG transcriptional data shared significant overlap of DEGs with previous studies of patient-derived iPSC cortical^42^ and neural cells^50^. Overlapping genes with the neural population highlight transcriptional regulation, development (particularly Hippo signaling, also in^114^), and synaptic signaling. Meanwhile, overlapping DEGs with the cortical population highlight cellular proliferation, development, and cellular survival. Developmental pathways (particularly WNT signaling) were also highlighted in our previous work looking at transcriptional and epigenetic (K4Me3 ChIP) dysregulation in HD iPSC striatal neurons^43^. That study identified dysfunction in cell cycle progression and cellular proliferation, which is which we find occurs not only because of CAG expansion but of HTT loss as well. Overlap between these two iPSC-derived populations and our *HTT* KO eCN transcriptional data was also significant (although only marginally for the cortical population), further highlighting the contribution of HTT loss of function to HD disease progression.

Interestingly, a subset of the DEGs appear consistent with changes we observed using our morphological feature analysis and may help unravel how the gene expression changes relate to cellular changes associated with gain or loss of function that occur because of mHTT. Harnessing the power of ML will greatly enhance the ability to unravel cellular signatures and mechanisms that occur in disease but has been underutilized by biologists. For large sophisticated datasets such as those that are acquired by imaging are perfectly suited to ML applications to find complex signals not readily detectable by conventional approaches^115^. A DNN was previously used to distinguish between CAG-expanded and *HTT* KO from controls in the RUES derivatives that contained densely packed clusters of cells. This finding suggested that there were predictive signals that set apart CAG expanded, *HTT* KO from control organoids. However, the study did not report the underlying signal nor investigated gene expression changes that could be the basis for the classifier they built^30^. For complex neural networks, there are methods to reveal the signals that drive classification decisions^115,116^. However, by using engineered features, like those used here, the discerning signals that separate groups become significantly more interpretable. Instead of relying on conventional machine learning approaches, we developed a novel method to track how features change over time, using the MCOT interaction term. This allows us to monitor dynamic cellular-based transformations that could reveal critical disease mechanisms, which might otherwise be overlooked by analyzing images at isolated time points. Similar work has been performed using pseudotime analyses of scRNA data to examine dynamic gene expression changes across time^117,118^. But, to our knowledge this has not been done with image data and therefore presents an innovative approach to study disease states.

For future work, one could interweave the MCOT with the OMICs signatures in order to gain deeper insights into which classes of genes drive cellular processes. For example, a subset of the morphological changes observed in the 72CAG and *HTT* KO eCNs parallel observed OMICs alterations. Size-related features such as soma and cell area (**Figure 2D, E Supplemental Figure 5)** showed significantly different MCOTs between 72CAG eCNs and controls. The cell area of 72CAG eCNs expanded more rapidly than that of the control eCNs, which remained relatively static. On the other hand, the cell area in *HTT* KO decreased compared to both 72CAG and control eCNs (**Figure 2D, E and Supplemental Figure 5A)**. These morphological changes align to changes in gene expression pathways altered such as “negative regulation of cell growth” which is downregulated in the *HTT* KO while upregulated in the 72CAG eCNs (**Figure 3C)**. In a similar but opposite fashion, expression changes are observed in the class of genes that make up “positive regulation of developmental growth” were upregulated in *HTT* KO whereas they were downregulated in the 72CAG eCNs (**Figure 3C)**. Variations in gene expression governing cell growth may account for the differences in size demonstrated here.

It’s also enticing to speculate that our novel feature– based paradigm captures gain and loss of function aspects of HTT. For example, the MCOT for some features were found to differ in unique ways between the control, *HTT* KO and 72CAG eCNs. The MCOT for a subset of the GLCM features “GLCM Disim, GLCM Corr, soma skew, FFT and “mean filter” was similar in control and 72CAG eCNs, while the MCOT for *HTT* KO eCNs differed drastically from control or 72CAG (**Figure 2 D, E and Supplemental Figure 5**). Genes in pathways uniquely dysregulated in *HTT* KO (**Figure 3, 5**) may impact cellular aspects altered due to the loss of HTT function that are entirely independent of the CAG repeat expansion.

The MCOT of other features (soma area, cell area, HOG, sholl Med, Mink Dim, sobel noise, sobel med) differ across all groups, but to different degrees and directions. These features may represent rate changes that reflect a combination of both loss and gain of function caused by the CAG repeat expansion. The DEGs that are found dysregulated in both the 72CAG and *HTT* KO eCNs may be drivers of these cellular structural changes (**Figure 3, 5**). In contrast, the MCOT for other features, such as “eccentricity” and “Frangi”, were found to be uniquely different when comparing the control to the 72CAG eCNs, but were not different between KO versus the control or 72CAG eCNs (**Figure 2D, E and Supplemental Figure 5**). These features may be attributes arising from the expansion of CAG repeats and may signify components of cellular gain-of-function.

Other morphological changes are consistent with gene expression changes. The edge feature captures both size and the complexity of an object. The edge MCOT was higher in the control group compared to 72CAG and *HTT* KO eCNs, indicating increased complexity in control eCNs (**Figure 2D,E).** The *HTT* KO eCNs exhibited greater dysregulation compared to the 72CAG eCNs. The direction of these changes match RNA expression levels of genes associated with “cell-cell contacts,” potentially influencing cellular complexity or size. For example, the morphology change was most pronounced in the *HTT* KO line compared to the 72CAG eCNs (**Figure 2E**) as is observed in the RNA expression (**Figure 3C).**

The majority of “complexity” features all decreased over time all groups, suggesting a decrease in cell-shape complexity as the neurons matured. However, the MCOT for complexity features decreased more in the control eCNs compared to the 72CAG and *HTT* KO eCNs (**Figure 2 D and Supplemental Figure 5 I, J, K, L)**. Gene expression changes observed in the class of genes that make up “actin cytoskeletal reorganization”, negative regulation of cell migration”, “actin filament organization” were all downregulated in the 72CAG and the *HTT* KO eCNs compared to controls (**Figure 3C).** Genes in these pathways drive the underlying forces of organization, structure, and cellular movement and since these changes are consistent with the observed morphological changes and may underlie developmental alterations in the 72CAG and *HTT* KO eCNs. Understanding how OMICs alterations caused by the loss of HTT or the presence of the CAG expansion relate to dynamic cellular characteristics could shed light to understanding the mechanisms by which gene expression drives pathological changes in HD. Previous studies have linked small-molecule induced transcriptomic perturbagens to cell morphology alterations^119,120^. Correlating gene-expression changes to feature-based morphological signatures may be able to help unravel the complexity and timing of when and how cellular systems fail in HD.

A recent study used a similar multi-omic network integration approach to identify links and shared processes between congenital heart disease and autism spectrum disorder^121,122^. Indeed, multi-omic network integration can generate richer hypotheses for assessing cause-and-effect relationships between disease-associated-omics and phenotypes of interest. To begin to assess how nodes in our network influence neurodegeneration, we annotated genes for whether they were essential in iPSC-derived cortical excitatory neurons (**Figure 6E**). This analysis identified *RFC2*, *THBS1*, *HAT1*, *PHF6*, *IPO7*, and *COG7* that are each down-regulated in CAG-expanded and *HTT* knockout eCNs and reduce survival upon CRISPRi knockdown (**Figure 6E**). This approach is similar to that employed by the NeuroLINCS Consortium^85^, where literature mining of ALS-associated genes and functional validation in *Drosophila* showed that nodes in disease-associated networks could be targets for *C9orf72*-mediated neurodegeneration^85^. Our analysis provides a platform for nominating pathways and genes that could be targets for therapeutic intervention in HD, and this type of analytical framework can be applied to biological processes involved in multiple neurodegenerative diseases. Future work can use these networks as the basis for implicating genes or proteins in influencing phenotypes such as neuronal survival or the modulation of a proposed biological pathway, nominating them as targets for genetic or pharmacological perturbations. Finally, the feature-based morphological analysis used here may reflect both gain and loss of function properties of the CAG expansions in HTT and could serve as a useful platform for future small molecule, drug or genetic screens that may correct the cellular consequences of mHTT.

## Supporting information

Supplemental Table 7 (eCN DEGs)

Supplemental Table 5 (ESC DEGs)

## Acknowledgements

This work was supported by grants from the CHDI Foundation, NIH/NINDS R01 NS089076 (E.F.), R35 NS116872 (L.M.T.), and R37 NS101996/NS/NINDS. Additional support was provided from NINDS/NIH T32 (NS082174 & NS121727-01, J.T.S.), and the Barbara Weedon Fellowship (M.J.L.). We thank Katerina Beck for technical assistance as well as Kelley Nelson, Gayane Abramova, Tami Tolpa, Katy Claiborn. We also thank the Genomics Research and Technology Hub Shared Resource of the Cancer Center Support Grant (CA-62203) at the University of California, Irvine for facilities and assistance in carrying out studies. The content is solely the responsibility of the authors and does not necessarily represent the official views of the National Institutes of Health.

## Author contributions

Conceptualization, J.T.S., M.J.L, Y-X. X, C.S., D.P.F., T.F.V., J.V.E, S.F.,J.A.K., E.F, L.M.T; Methodology, J.T.S., M.J.L., K.Q.W., C.S., J.A.K., Z.F, C.P, R.T, S.L., N.A,; Software, M.J.L, Y.R., J.W., R.M., S.L, R.T, N.A,; Formal Analysis, J.T.S., M.J.L, Y-X. X., O.W., Y.R., J.W., R.M., J.A.K. R.T, S.L, M.S, Z.F, K.S; Investigation, J.T.S., Y-X. X., K.Q.W., J.A.K., O.W., S.L, Z.F, K.S, H.J, T.H, M.F, C.P, L.E, E.D, Resources, D.F.; Data Curation, J.T.S., M.J.L, Y.R., K.W., S.L, N.A,; Writing – Original Draft, J.T.S., M.J.L., Y-X. X., J.A.K., L.M.T.; Writing – Review & Editing, J.T.S, M.J.L., Y-X. X., D.P.F., T.F.V., C.S., S.F., J.V.E., J.A.K., E.F., L.M.T.; Visualization, J.T.S., M.J.L., R.M., S.L, J.A.K.; Supervision, J.V.E., J.A.K., S.F., E.F., L.M.T.; Project Administration, J.V.E., S.F., L.M.T., E.F., J.A.K.; Funding Acquisition, E.F., L.M.T., S.M.F, J.A.K

## Supplemental information titles and legends

### Supplemental information

Document S1. Figures S1–S11 and Table S1-S4, S6

Table S5, S7-S8. Excel files containing additional data too large to fit in a PDF

## Methods

### Pluripotent stem cell culture and sample generation

Isogenic human pluripotent stem cells derived from the RUES2 (<u>NIHhESC-09-0013)</u> embryonic stem cell line were acquired from Coriell through the CHDI foundation. Lines were maintained on Geltrex basement membrane (Thermo Fisher # A1413301) in mTESR1 medium (Stem Cell Technologies #85850) for omics studies and on Growth Factor Reduced (GFR) Matrigel (Corning # 356231) and mTESR plus medium (StemCell Technologies,# 05825). Cells were grown at 37°C and 5% CO2 and were passaged using ReLeSR (Stem Cell Technologies) when they reached 70-80% confluency. Colonies were cryopreserved in CryoStor CS10 (Stem Cell Technologies). Cell pellets for each of the Omics assays were generated and harvested in parallel for all cell lines, from 3 successive passages using the same lot of each reagent.

### Cortical Neuron differentiation and sample generation

#### Omics Studies

Pluripotent cells were dissociated to single cells with Accutase (Fisher Scientific # NC9839010) and replated in mTeSR1 plus rock inhibitor (RI) (Peprotech # 1293823) at a density of 4×10^6^ cells in laminin (Biolamina #LN521) coated T25 flasks. T25 flasks were fed mTeSR1 for no more than 2 days until they reached >80% confluence at which point mTeSR was replaced with CM1_o_ consisting of Base C media plus 10 µM SB 431542 (Stem Cell Technologies #17502048), 200nM LDN193189 (Stem Cell Technologies, #72147) and 5 µM Cyclopamine (Stem Cell Technologies #72074). Base C media consists of 0.5X DMEM/F12 (Thermo Fisher, #10565042), 0.5X Neuraobasal (Thermo Fisher #21103049), 0.5% N2 (Thermo Fisher # 17502048) and 1% B27(Thermo Fisher #17504-044). From this timepoint (day 0), the cells were fed daily with 10 ml of CM1 until day 8. At d8, cells were passaged 1:2 using cell scraping and gentile trituration onto Poly-ornithine (Sigma # P4957) and 20 ug/mL laminin (Sigma Aldrich #L2020) coated T25 flasks. The cells (at this stage, referred to as NPCs) were then fed everyday with CM2_o_ composed Base C media plus 2.5ug/ml Insulin (Thermo Fisher #12585-014), 0.5% NEAA (Thermo Fisher #11140050), 5 uM Cyclopamine and 20ng/mL FGF-2 (Peprotech #100-18B). On d15, neural progenitors were dissociated using Accutase and banked in CryoStor CS10. D15 progenitors were banked from 2 rounds of successive passages at the pluripotent stage for each line using identical lots of materials (4-6 batch replicates per line).

Banked progenitors for all lines were thawed in parallel, one batch at a time (4 batches per line), onto Poly-ornithine/laminin 6 well plates at a density of 4×10^6^ cells/well in CM3_o_ comprised of Base media C plus 2.5ug/ml Insulin (Thermo Fisher #12585-014), 0.5% NEAA (Thermo Fisher #11140050) plus 2 uM Cyclopamine, 10 ng/ml BDNF (Peprotech #450-02), and 10 ng/ml GDNF (Peprotech # 450-10).). At this point, the cells were considered early-stage cortical neurons (es-CNs) and were fed every day for 6 days. At d21, cells were dissociated using Accutase and replated for final maturation onto Poly-D-lysine (Sigma Aldrich #P6407) coated 12 well plates, at a density of 2-2.5 x10^6^ cells per well. D21 eCNs were fed every other day for 2 weeks in CM4_o_ comprised of CM3_o_ supplemented with 10 µM DAPT, 20 ng/mL BDNF, and 20 ng/mL GDNF. At day35, eCNs were washed 3x with PBS (Thermo Fisher # 10010023), harvested using a cell scraper, and centrifuged to pellet. Transcriptomics and proteomics pellets were flash-frozen dry, epigenetics pellets (ATACseq and ChIPseq) were resuspended in CryoStore CS10 and frozen at −80.

#### Imaging studies

Pluripotent cells were dissociated to single cells with Accutase (Fisher Scientific # NC9839010) and replated in mTeSR Plus with rock inhibitor (RI) (Selleck Chemicals, #S1049) at a density of 3×10^6^ cells per GFR-coated T25 flask. T25 flasks were fed mTeSR Plus for no more than 2 days until they reached >90% confluence, at which point mTeSR was replaced with CM1i consisting of Base C media plus 10 µM SB 431542 (Tocris, #1614), 400 µM LDN193189 (Tocris, #605) and 1 µM cyclopamine (Tocris #1623). From this timepoint (day 0), the cells were fed daily with 10 ml of CM1 until day 8. At d8, cells were passaged 1:2 using Accutase and plated onto onto Poly-ornithine (Sigma # P4957) and 20 ug/mL laminin (Sigma Aldrich #L2020)-coated T25 flasks. The cells (at this stage, referred to as NPCs) were then fed every other day with CM2_i_ until day 15. CM2 consists of: Base C media plus MEM non-essential Amino Acids (Life Technologies, #11140050), 0.1 mM 2-mercaptoethanol, 2.5 ug/ ml insulin, 1 µM cyclopamine and 20 ng/ml FGF. On d15, the NPCs were disassociated using Accutase, and frozen at 8 million cells per vial in Stem Cell Banker cryopreservation solution (CedarLane Labs, # 11890). For each experiment, a vial of each cell line was thawed and plated in CM3 media into a 6-well plate pre-coated with Poly-L-Ornithine and 20 ug/ul of laminin at a density of 4 million cells per well. CM3_i_ media consists of Base C media plus MEM non-essential amino acids, 0.1 mM 2-mercaptoethanol, 2.5 ug/ ml insulin, 1 µM cyclopamine, 10ng/ml BDNF (R&D Systems, #248-BD) and 10ng/ ml GDNF (R&D Systems, #212-GD. At this point, the cells are considered es-CNs. The es-CNs were fed every day for 6 days until day 21. On day 21, e-CNs were passaged to 384 plates that were precoated with poly-L-ornithine, laminin and fibronectin (CB40008A) in CM4 media. CM4_i_ media consists of 0.5X DMEM/F12, 0.5X Neurobasal, 1X MACS NeuroBrew-21 Supplement (Myltenyi, # 130-093-566),1X N2 Supplement, 2.5 ug/ml insulin rh, zinc solution, 10 µM DAPT (R&D Systems, #2634), 20 ng/ml BDNF, 20 ng/ml GDNF and 1X Anti-Anti. After day differentiation day 27, these cells are considered late-stage cortical neurons (eCNs).

### Immunocytochemistry and microscopy—omics studies

Cells were fixed with 4% paraformaldehyde (Fisher Scientific # 50980487) for 10 minutes at room temperature, then washed three times with PBS (Corning # 21030CV). Cells were permeabilized with 0.3% Triton-X (Sigma #T8787) in PBS for 10 minutes and then blocked with 2% goat serum (Thermo Fisher #16210-064), 3% BSA (Thermo Fisher # 15260-037), 0.1% Triton-X, and 0.3M Glycine (Fisher # BP381-1) in PBS for 1 hour at room temperature and then incubated in primary antibody diluted in block, overnight at 4°C (anti-SOX2 (1:500) Millipore AB5603, anti-OCT4 (1:300) GeneTex GTX627419, anti-PAX6 (1:250) AbCam ab5790, anti-Nestin (1:1000) Millipore MAB5326, anti-Tubuin, B III (1:1000) Millipore MAB5564, anti-FOXG1 (1:300) Abcam ab18259, anti-CTIP2 (1:500) Abcam ab18465, anti-MAP2 (1:1000) Synaptic Systems 188004, anti-TBR1 (1:250) Abcam ab31940). Primary antibody was removed, and cells washed three times with PBS and then incubated for 1 hour in secondary antibody diluted 1:1000 in block, in the dark at room temperature (Alexa Fluor Goat IgG (H+L) Secondary Antibody, Thermo Fisher Scientific). Cells were washed with PBS for three times and then washed in PBS containing Hoechst 33342 (Sigma #14533) for 10 minutes and then a final wash in PBS. Coverslips were mounted with Fluoromount-G® (Fisher # OB10001) and allowed to dry. Images were acquired on a Keyence BZ-X810 Widefield Microscope. Cell populations were quantified using Cell Profiler (https://cellprofiler.org/) using pipelines modified from those described^66^ with n=3-4 images across each differentiation replicate per line.

### Immunocytochemistry—Imaging studies

eCNs were fixed at day ∼35 by permeabilization using a 0.1% Triton-X/PBS solution for 20 minutes at room temperature. Permeabilization solution was removed and a 1 M glycine solution added and incubated at room temperature for 20 minutes. A blocking solution of 0.1% Triton-X/PBS, 2% FBS, and 3% BSA was added after removal of the glycine solution and incubated at room temperature for 1.5 hours. Antibodies against: **MAP2** (1:1000 Abcam chicken-anti MAP2 # ab5392), **KI67** (1:200 Millipore mouse-anti KI67 # mab4190), **CTIP2** (1:250, Abcam, #ab 18645), **TBR1** (1:2000, Abcam Ab183032), **FOXG1** (1:2000, Abcam #ab18259), and DARPP-32 (1:250 abcam, # ab40801) were diluted in blocking solution and incubated overnight at 4°C. Primary antibody was removed by washing cells 3 times with 0.1%Triton-X/PBS. Secondary antibodies all from Invitrogen (goat anti chk 647, #A21449; donkey anti-Rb 488, #A32790; goat anti-mouse 488; #A11001 or goat Anti-Rat 488, #A11006) were added to 1:1000 in blocking solution and incubated at room temperature and covered for 1.5 hours. Cells were spun down at 8000 RPM in a cold centrifuge during the washes. Cells were washed in PBS for 5 minutes, then once with PBS plus Hoechst at a dilution of 1:1000 and incubated for 10 minutes at room temperature, covered. The Hoechst was washed out with PBS and the cells covered in PBS for imaging.

#### CellProfiler

To measure the patterning propensity for the eCNs, cells were fixed, stained with various antibodies as described above, and images were subjected to a modified CellProfiler^124^ pipeline to examine the percent of cells staining for a particular antibody, similar to the colocalization pipeline as described here (https://cellprofiler.org/examples). Images were pooled in CellProfiler and analysed on a per-image basis. Background values were calculated per image then subtracted from the whole image. Following background correction, all objects in each DAPI image were identified as nuclei using minimum and maximum diameters per object and filtering out excessive intensity values using a minimum cross-entropy thresholding method^124^. This method identifies all possible nuclear objects within the appropriate size range. Intensity values were calculated per object then used to filter out non-nuclear objects or dead cells that display bright nuclei, and then relabelled as nuclei segments. Images that contained KI67/ TBR1/ FOXG1/CTIP2 or DARPP32 staining were subjected to a feature enhancement step that increases the signal-to-noise ratio. Enhanced images were run through segmentation to identify all objects considered “positive” in terms of size and signal intensity again using a minimum cross-entropy thresholding method^124^. These identified objects were renamed as “positively labelled” segments. These segments were related to the nuclei segments to determine how many nuclei were “positive” for that particular antibody. Finally, the number of “positively labelled” cells were divided by the number of total nuclei to calculate the percent of KI67/ TBR1/ FOXG1/CTIP2 or DARPP32 “positive” cells. The segmentation overlays and math were exported from the program and then plotted in Prism10^125^.

Cell Profiler was also used to measure neurite length was based on Tian *et al.* 2019^66^, with some additional modifications to the workflow as previously described^26^. The same background correction and primary object identification as the analysis described above was performed but on images from differentiation day ∼32 from live cells that express SYN:EGFP. The soma for each neuron was identified in CellProfiler^124^ and segmented based on size, eccentricity, and intensity of SYN:EGFP signal. Soma segments were filtered by eccentricity to eliminate any potentially dead, rounded cells or debris from the analysis. Next, the images were run through feature enhancement to increase contrast between potential neurites and the background. Neurites were identified using the filtered soma masks as points of origin for neurite outgrowth and an Otsu thresholding method^124^. Skeleton length as well as trunk and branch end numbers were calculated and exported as a spreadsheet. We normalized each experiment to account for variation across batches of differentiated cells, as previously described^50^. Briefly, we calculated the average neurite length of control eCNs for each batch and normalized each individual value by this average to generate an averaged ratio for each neurite measured. These ratios are used in the histogram in Figure 2.

#### Statistical analysis

We developed a statistical package called RMeDPower^123^ in R, a complete package of statistical tools that allow a scientist to understand the effect size and variance contribution of a set of variables within a dataset to a given response^123^. RMeDPower uses linear mixed models on repeated measures data such as those described here. Outliers were removed using the Cooks distance^126^ and data was log transformed for statistical analysis. All p values and estimates for each comparison for the neurite analysis and percent positive staining were calculated in RMeDPower^123^.

### RNAseq

RNA was processed as previously described^85^ from the 15 pluripotent samples (5 lines in triplicate) and 16 cortical neuron samples (3 replicates for all lines except *HTT*KO which had 4 replicates as the 3^rd^ replicate for proteomics and epigenomics assays were derived from separated differentiation batches due to insufficient yield). Briefly, the RNeasy mini kit was used along with QIAshredders for homogenization and DNase I for gDNA elimination (Qiagen #74106, # 79656, #79254). The RNA samples were then analyzed for an RNA integrity number (RIN) which was determined to be greater than 9 for each sample except for one, which had RIN of 7.3 (which later passed QC at the library stage). rRNAs were removed and libraries generated using TruSeq Stranded Total RNA library prep kit with Ribo-Zero (Qiagen). RNA-seq libraries were titrated by qPCR (Kapa), normalized according to size (Agilent Bioanalyzer 2100) with sufficient paired end (PE) sequencing cycles to obtain 50 million PE reads per sample.

#### RNAseq Data Analysis

Reads were mapped to the GRCh38 reference genome, QCed, and gene expression and differential expression were quantified using Hisat2, featureCounts^127^ (PMID: 25260700) and DESeq2 (PMID: 25516281). Normalized and transformed count data were then used for exploratory analysis and differentially expressed genes (DEG, FDR <0.1) were analyzed with commercial and open-source pathway and network analysis tools, Ingenuity Pathway Analysis (IPA) and Gorilla gene ontology (GO) analysis.

### Epigenetic Data

#### ATAC-seq

We used 50,000 nuclei for the transposase reaction, following the method described previously^128^. Subsequently, samples were purified with the DNA Clean & Concentrator–5 Kit (Zymo Research), and then PCR-amplified using Nextera indexing primers (Illumina). The number of PCR cycles were optimized by qPCR to avoid over amplification of the libraries. The final libraries were purified using 1x PCRClean DX beads (Aline Biosciences). The enrichment of accessible chromatin regions in the libraries were assessed by qPCR using primers mapped to open chromatin region and gene desert regions^129^. The libraries fragment distribution was analyzed by a Fragment Analyzer™ instrument (Advanced Analytical), and the concentration measured by a qPCR-based method (KAPA Library Quantification Kit for Illumina Sequencing Platforms). PE sequencing (40 nt) was conducted using Illumina NextSeq platform at the MIT BioMicroCenter. The quality of the data was assessed using FastQC and the sequences aligned to the HG38 reference genome build using BWA, a short sequence aligner, with default parameters. Aligned reads with a mapping quality of less than 10 and mitochondrial reads were removed using SAMtools. Duplicate reads were removed using PicardTools.

#### ChIP-seq

Histone modification ChIP experiments were performed using antibodies against H3K27me3 (C36B11, Cell Signaling), H3K4me1 (Abcam # ab8895,), H3K4me3 (Millipore #07-473), H3K27ac (Cell Signaling # D5E4) and IgG. For each ChIP, 1X10^6^ cells were incubated in lysis buffer (50 mM Tris-HCL pH 8.0, 150 mM NaCl, 1% Triton X-100, 0.1% Na-deoxycholate, and 5 mM CaCl2 supplemented with protease inhibitors and 10 mM sodium butyrate for K27ac ChIP) for 20 min on ice. The chromatin was digested to 1-5 nucleosomes using Micrococcal Nuclease (NEB # M0247S), at concentration determined by titration experiments, for 10 min at 37°C. To terminate the digestion, 20 mM EDTA was added to the samples. The antibody was added and incubated overnight at 4°C. On the next day, 25 µl of protein G Dynabeads (Thermo Scientific # 10004D,) was added and the samples rotated for 2 hr at 4°C. The beads were washed 6 times: 2 washes with RIPA buffer (10 mM Tris-HCl pH8, 1 mM EDTA, 140 mM NaCl, 1% Triton X-100, 0.1% Na-deoxycholate, 0.1% SDS), 2 washes with high salt buffer (10 mM Tris-HCl pH8, 1 mM EDTA, 360 mM NaCl, 1% Triton X-100, 0.1% Na-deoxycholate, 0.1% SDS), 2X washes with LiCl buffer (10 mM Tris-HCl pH8, 1 mM EDTA, 250 mM LiCl, 0.5% IGEPAL CA-630, 0.5% Na-deoxycholate) and 1 wash with TE (10 mM Tris-HCl pH8, 1 mM EDTA). The DNA was eluted by incubation of the beads in elution buffer (10 mM Tris-HCl pH8, 5 mM EDTA, 300 mM NaCl, 0.1% SDS, 50 µg Proteinase K) for 1 hr. at 62 and purified with 1X PCRClean DX beads (Aline Biosciences). The sequencing libraries were constructed using the NEBNext^®^Ultra™ II DNA Library Prep Kit for Illumina (NEB # E7645S), followed by analysis using Fragment Analyzer™ instrument (Advanced Analytical), and quantification by a qPCR-based method (KAPA Library Quantification Kit for Illumina Sequencing Platforms). Single-end sequencing (50 nt) was conducted using Illumina HiSeq 2000 or NovaSeq S4 platform at the MIT BioMicroCenter. QC of the sequences, genome alignment and removal of low-quality reads and duplicates were done as described for ATAC-seq. Regions marked by the specific histone modification (peaks) with MACS2 were determined using the “broad peak” setting, a “broad” cutoff of 0.1, a q-value cutoff of 0.05, and the aligned reads from an IgG ChIP sample as a control.

#### Processing epigenetic data

We processed the ATAC-seq data using the ENCODE-DCC ATAC-seq pipeline v.1.7.1. Using this pipeline, we identified FASTQ read adaptors with the detect_adapter.py script in GGR_code (https://github.com/nboley/GGR_code). We trimmed read adaptors with the trim_adaptors function in cutadapt 1.9.1. Processed FASTQs, were aligned using Bowtie2 version 2.2.6, requiring SAMtools version 1.7 and SAMstats version 0.2.1. Picard tools version 1.126 MarkDuplicates was used to deduplicate the aligned reads. These tools were used with default parameters except for setting the parameters “atac.auto_adect_adapter” and “atac.enable_xcor” to “true”.

The ChIP-seq data was analyzed using the ENCODE-DCC ChIP-seq pipeline v1.3.6. In this pipeline, FASTQ data were cropped with Trimmomatic 0.39, aligned with Bowtie2 version 2.26 and deduplicated with Picard version 1.126 MarkDuplicates. Default parameters were used for all arguments except for setting “chip.pipeline_type” to “histone”.

#### Analysis of epigenetic data

From the aligned, deduplicated ATAC-seq data, MACS2 version 2.1.0 was used to identify peaks that had significant local enrichments at a p-value threshold of 0.01. For the aligned, deduplicated ChIP-seq data, peak calling was performed using SPP version 1.14, GEM 2.4.1 and PeakSeq version 1.25. IDR 2.0.4 was run to determine a high-quality set of consensus peaks from the results of the multiple peak callers. Upon identifying peaks, DiffBind version 2.10.0 was used to perform differential analysis between ChIP-seq or ATAC-seq peaks for the following comparisons: *HTT* knock-out vs. control, *HTT* HD knockdown vs *HTT* wild-type HD, 56/22 (adult onset) HD vs control, and 72/20 (expanded repeat length) HD vs control. We refer to the union of peaks found in 20/20 and 22/20 cells as “control” in these comparisons. DiffBind in R version 3.5.2 was used to identify differential peaks as those with an FDR-adjusted p-value less than 0.1

To determine similarities and differences across cortical neuron replicates and genotypes in the ATAC-seq or ChIP-seq data, Principal Component Analysis (PCA) was used as implemented in Scikit-Learn version 0.23.2. To preprocess the data, peaks were limited to those that were within 5kb from a protein-coding gene. The signal of the filtered peaks was standardized before performing PCA with 10 principal components. Replicates were removed from the analysis if they failed the ATAC-seq or ChIP-seq ENCODE benchmarks determined from FastQC (http://www.bioinformatics.babraham.ac.uk/projects/fastqc/).

We determined enriched, known motifs from the ATAC-seq and ChIP-seq marks by applying HOMER version 4.11 to the significantly differential ATAC-seq or ChIP-seq peaks inferred from DiffBind (http://bioconductor.org/packages/release/bioc/html/DiffBind.html). The background for the motif enrichment analysis was the set of all peaks identified by MACS2 for a given genotype, across replicates. For motif enrichment, we looked for motifs in 200 base-pair regions in repeat-masked sequences.

### Proteomics

Upon receipt, 21 cortical neuron samples from 7 lines (3 replicates of each) were lyophilized and lysed in Tris-Urea lysis buffer (33mM Tris, 1.33mM CaCl2, 4M urea, 0.67mM DTT in 1M NH4CO3, pH 8.3-8.7) with sonication. Protein BCA assay was performed, and a volume required for sufficient protein was taken from the cell lysate for the proteomic sample preparation on Beckman Biomek i7 Automated Workstation. The samples were reduced with 50mM TCEP, alkylated with 200mM iodoacetamide (IAA), digested by Trypsin/LysC, followed by the peptide desalting by using Waters Oasis HLB plate. Desalted tryptic peptides were dried down in a SpeedVac and reconstituted with 0.1% aqueous formic acid for data independent acquisition (DIA)-mass spectrometry (MS) analysis.

#### LC-MS Analysis

The peptide samples were separated on a 45-minute gradient, a window of 15 m/z,on the UltiMate 3000 HPLC system (Thermo Scientific) connected with Orbitrap Fusion Lumos Tribrid mass spectrometer (Thermo Scientific). Here, mobile phase A and B consisted of 0.1% formic acid in water and 0.1% formic acid in 100% acetonitrile, respectively, using a gradient of 6-36% B in 41 min, 36-80% B in 1 min, and 80% B in 1 min at the flow rate of 9.5 ul/min. Resolved peptides were ionized by an EASY-Spray ion source. In MS scans, internal mass calibration was performed using EASY-IC. Mass spectra were acquired in a data-independent manner, parameters for the DIA method are as follows: resolution at 60,000, mass range of 400-1000 m/z, and maximum injection time of 50 ms for MS1 scan; resolution at 15,000, HCD collision energy of 30%, mass range of 400-1000 m/z, and maximum injection time of 30 ms for MS2 scan. For both MS1 and MS2, RF Lens 30% and normalized AGC Target 150% were applied.

#### Proteomics Data Analysis

MS data files were analyzed in DIA-NN v.1.8 (Demichev et al, 2020), using the spectral library-free search feature with Prosit against the Human Uniprot database. For cohort comparisons of cortical neuron samples, the DIA-NN generated protein intensities were log base 2-transformed and standardized across each sample so that the protein quantity distribution in each sample had a mean of zero and standard deviation of one. Proteins quantified in every sample were compared between the control groups and HD or KO groups respectively using t-tests with Benjamini-Hochberg correction.

### Longitudinal single cell imaging and analysis

eCNs were transduced with pHR-hSyn:EGFP (Addgene #114215)^130^ or GEDI^131^ at ∼5 MOI after plating in 384-well format. eCNs were subjected to RM and imaged daily as previously described^25,52–57,132–135^ using ImageXpress Micro Confocal High-Content Imaging System from Molecular Devices for 7-10 days starting at differentiation day ∼28-32. Images of different microscope fields from the same well were stitched together into montages, and montages of the same well collected at different time points were organized into composite files in temporal order. Image analysis was performed in a computational pipeline constructed within the open-source program Galaxy, to identify individual cells to perform survival analysis, or apply morphological measurements as described below. eCNs were hand tracked using previously described methods and a Cox mixed effects proportional hazards analysis^61^. We compared the survival to the group of 20CAG lines (20CAGn3 and 20CAG44) to the 72CAG lines (72CAGn2, 72CAGn3 and 72CAGn1). We also compared the survival rates of the control compared to the KO eCNs.

#### Feature extraction

After eCNs were subjected to RM as described above, images from control 20CAG, expanded 72CAG and KO lines were processed in our custom-built image-processing pipeline Galaxy software^71^. Images from the same well were stitched together into montages, and cells were identified and segmented using a thresholding method based on the standard deviation from the mean to ensure that brighter cells are accurately identified and isolated. Images were subjected to our custom-built filtering algorithm written in Python to crop each detected object into 3003×00 pixel size and remove any crop with edge artifacts, dead cells, clumps, out-of-focus, images with bad intensity signal, or non-centered cells to ensure cell quality for further analysis. These are called “cell crops”. This was achieved through a series of well-defined steps. First, cells located on the image edge that could not be cropped into 300×300 pixel region were excluded. The remaining cells were cropped and subjected to a Gaussian blur algorithm^136^ to smooth the image and reduce high-frequency noise to feed into Otsu thresholding^137^, which separates objects from background. These segmented objects were then subjected to the Erosion and Dilation algorithm^138^, which acts to shrink and enlarge image foreground to detect and separate adjacent cells and eliminate small artifacts. Images with more than one potential cell were filtered out to ensure our analysis only focused on individual cell images for single cell analysis. In addition, cells that fell outside the defined range of minimum and maximum area were also filtered out. This helps to exclude cells that were either too small to be of interest, often indicating artifacts or dead cells, or too large, suggesting the presence of clumps. The cell crops were then fed into our custom-made feature extraction algorithm. The feature extraction algorithm is written in Python and contained within a Jupyter notebook (https://github.com/finkbeiner-lab/FeatureAnalysis)-Morphological properties such as cell area, cell texture, complexity and intensity were analyzed by the following steps. Various features such as median value, and distribution skewness statistics were extracted using a combination of image processing packages, notably OpenCV^31^, NumPy^32^, Scikit-Image^33^, and SciPy^34^ library. To obtain these features, a median blur filter^139^ preprocessing was applied to the image crops. Subsequently, a binary threshold process was employed to selectively capture only the soma (the cell body of each neuron) for analysis. This generates binary masks which were then used to calculate key statistics of each soma determined by the total pixel count within the mask; the Soma Median, representing the median pixel value of the soma area; and the Soma Skewness, which measures the asymmetric in the distribution value within the soma area. Additionally, cell area was extracted, which is the total pixel count of both the soma and neurite areas, to capture valuable information about the entire cell structure. A mean blur filter^139^ feature was used to smooth the image to reduce unnecessary noise. The median of the local gray-level histogram from the mean filter was calculated to provide insights into the image intensity of each cell line.

To analyze the shape and the morphology of the cells, edge detection methods were used to try to identify any distinct edge characteristics, such as Sobel operator^35^, HOG (histogram of gradients)^36^, Canny edge detection^37^ and Fast Fourier Transform (FFT)^38^. The Sobel operator^35^ was used to capture the mean, the median value, and the noise in the image, which highlight the areas of significant intensity changes in the cell image and are indicative of the presence of edges. HOG^36^, on the other hand, captures more information within the cells and more distinctive local patterns. Canny edge detection was used to capture sharp and continuous edges in the cell, and a High-Pass Filter (HPF) using Fast Fourier Transform (FFT)^38^ were applied to enhance high-frequency components, including edges and texture patterns.

To quantify the complexity in the cell, Minkowski-Bouligand Fractal Dimension^81^ feature was used to measure the roughness or complexity of an object. There are also two texture features including Haralick^140^ and GLCM(Gray-Level Co-occurrence Matrix)^79^ features that quantify the spatial relationships between pixel intensities in an image. The GLCM Correlation measures the probability occurrence of neighboring pixels in the horizontal and vertical directions, the GLCM Energy is the sum of the squared elements and captures the uniformity. The GCLM Correlation measures the probability of the neighboring pixel pairs ^141,142^.

The GLCM measures subtle texture differences between cell lines that may not be apparent through other features. To analyze the branching patterns, features such as Frangi Filter^143^, a vessel enhancement filter, and Sholl Intersections^84^ were used. Both can provide valuable information about neurites and the pattern of neurite outgrowth.

Values from all the features were extracted from the first (T1) and last timepoint (∼T7) available were saved into a CSV file for further downstream analysis. The multiple-timepoint data was combined and processed through a RMedPower^144^ tool kit that allows a user to examine the normality assumptions for the linear mixed model^144^ and adjust if necessary as well as using the Rosner’s test^145^ to remove any detected outliers on a per-feature basis. All of the feature values except for “edge” and “sholl med” were not normally distributed so they were log transformed as described^144^. To estimate this shift modulated by disease status we use a Linear Mixed Effects Model (LMM)^77,78^ called RMedPower^144^ to capture the mean feature values in control 20CAG, expanded 72CAG and *HTT*KO lines at each distinct timepoint and the additional shift across the two time-points due to a change in disease status via an interaction term. Hence, the interaction term is between the time and the disease status variables. The mathematical formulation of this LMM is described next.

Let 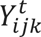 denote the feature value in the natural or the logarithm scale (see above) of the 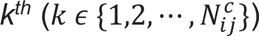 cell at time-point *t* (*t ϵ* {*T*1, *T*7}), derived from the *j^th^* (*j ϵ* {1,2, ⋯, *N*^*c*^}) cell-line assayed in the *i^th^* (*i ϵ Exp* = {1,2, ⋯, *N*^*e*^}) experimental batch. *N*^*e*^ is the number of experimental batches. *N*^*c*^ (5) is the number of cell-lines – 2 20CAG lines, 3 72CAG lines and 1 HTTKO line. 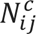 is the number of cells from cell-line *j* assayed in experimental batch *i*. The notation *A* and *R* refers to the sets of all alternate and reference cell-lines respectively that are used in the experiment. Then the LMM states,

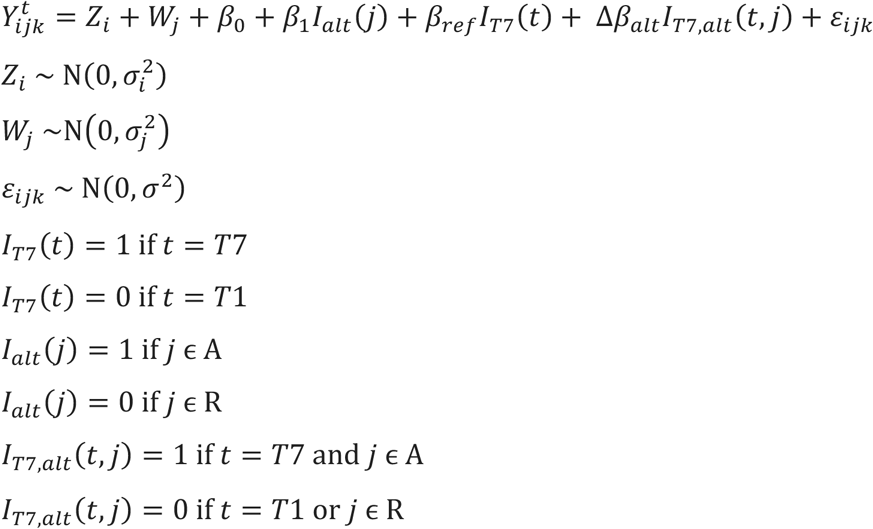

Where *Z*_*i*_ denotes the random effect associated with the *i^th^* experimental batch, *W_j_* denotes the random effect associated with the *j^th^* cell-line. Both these random effects are assumed to drawn from normal distributions with zero means and variances 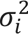 and 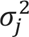 capturing the inter-experimental batch variances and inter-cell-line variances, respectively. The ɛ_*ijk*_ term captures the residual error of the model. Let *β*_*ref*_ refer to the change in the marginal expectation of the feature value over time for the reference cell-lines (e.g., 20CAG), i.e.,

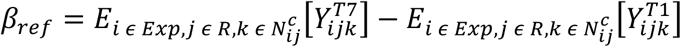

The *β*_0_ term captures the mean feature value at time T1 for the reference cell-lines, i.e.,

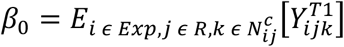

The *β*_1_ term captures the change in mean feature value at time T1 between the alternate and the reference cell-lines, i.e.,

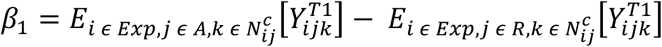

The subscripts for the expectation function, *E* indicates that the mean is taken by averaging out effects over all experimental batches, over all reference cell-lines and all cells in each of these reference cell-lines. *I*_*T*7_(*t*) and *I*_*T*7,*alt*_(*t*, *j*) are indicator functions. *β*_*alt*_ can analogously defined for the alternate cell-line (e.g., 72CAG) though is not directly estimated as one of the coefficients in the above model. Δ*β*_*alt*_ estimated by the coefficient of the interaction term in the LMM described above, captures the additional change over time of the mean feature value for the alternate cell-line. We call this interaction term the “Morphology Change Over Time” or “MCOT”(the time:disease_status interaction terms when scaled by the change over time in the 72CAG and *HTT*KO or control and expressed as percentages (**Figure 2E**). Specifically, MCOT is defined as,

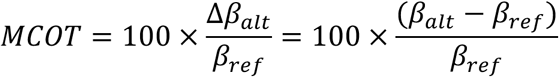

Significant differences in the rate of change of these features were identified by the significance of the interaction term. The LMMs were fitted using the lmerTest^77,78^.

Use of RM allows us to acquire very large sample sizes and therefore resulted in many observations, we observed very low p-values when comparing most all of the features, but in some cases this change was very small therefore, we considered the effect sizes in highlighting significant associations^59^ and only considered a change of 15% or greater to be considered a *bona fide* difference across the groups.

#### Identifying concordant changes in differential omics data

To define differentially expressed genes in the RNA-seq data, we used an FDR-adjusted p-value cutoff of 0.05 and an absolute log_2_ fold change cut off of 1. For the epigenetic data, we used an FDR-adjusted p-value cutoff of 0.1. For the proteomics, we used a nominal p-value cut off of 0.05 and an absolute log_2_ fold change cutoff of 0.6. To identify enriched pathways, we ranked each omics data by their log_2_ fold change and applied Gene Set Enrichment Analysis (GSEA) for each assay. As a reference pathway set, we used the 2021 Gene Ontology Biological Processes, as obtained from Enrichr at https://maayanlab.cloud/Enrichr/#libraries. To apply GSEA to proteomic data, we limited the background set of genes to proteins that were detected in the proteomics analysis to account for the observation that not every gene in the genome is detectable by mass spectrometry.

For each assay, we defined “concordant” changes as a differentially altered gene or protein in at least one of the HD or KO comparisons that had the same fold change direction in all three genotypes. For example, if gene A were differentially expressed in the RNA-seq data when comparing *HTT*KO to control, we would define the change as concordant if gene A was downregulated when comparing *HTT*KO to control, 56CAG to control and 72CAGn1 to control (20CAGn1 and 20CAGn2). This analysis was performed separately for each assay. To assess whether the set of differentially expressed genes or differentially abundant proteins were enriched for genes or proteins with concordant changes, we performed a hypergeometric test, where the background was either the total number of genes that were expressed in the cortical neuron data or the total number of proteins detected. Visualizations were performed in R using the packages ComplexHeatmap version 2.4.3 and UpSetR version 1.4.0. For GSEA, we used the R package fgsea version 1.14.0.

### Multi-omic integration analysis for HTT KO and HD cell lines

To identify the shared and distinct biological processes associated with *HTT* loss-of-function and HD. We utilized the Prize-Collecting Steiner Forest algorithm (PCSF) as implemented in OmicsIntegrator 2 (v2.3.10, https://github.com/fraenkel-lab/OmicsIntegrator2^89^). The PCSF algorithm identifies genotype-associated networks based on retrieving significantly altered omics signals without including low-confidence edges. The result includes the differentially altered omics and predicted interactors from the reference interactome, which we call “predicted nodes”. We used OmicsIntegrator to map proteomic, transcriptomic, and epigenetic motifs to a set of known protein-protein and protein-metabolite interactions derived from physical protein-protein interactions from iRefIndex versions 17 and 14 as well as and protein-metabolite interactions described in the HMDB and Recon 2 databases.

We performed differential analyses comparing *HTT* knock-out to the combined controls, 56/22 HD to the controls, and 72/20 HD to the controls. Differentially expressed genes were determined as those with an FDR-adjusted p-value less than 0.1 and absolute log_2_ fold change greater than 1 after applying DESeq2 to the transcriptomics data^146^. Differential proteomics were those with an FDR-adjusted p-value less than 0.1 determined by a t-test after normalization. Enriched motifs were determined as those with an FDR-adjusted p-value after motif enrichment analysis using HOMER^147^. To set the weights, or prizes, of the termini, we associated each differentially expressed gene, differentially abundant protein or enriched epigenetic motif with its negative log_10_, FDR-adjusted p-value. Prize values were minimum-maximum scaled by data type. For interpretability, we kept a proportion of the most enriched prizes for network integration; for the *HTT* KO network, the top 30 transcriptomic, proteomic and epigenomic prizes were included, and for the HD network the top 50 transcriptomic, proteomic and epigenomic prizes were selected from the 56/22 and 72/20 prizes. For the HD network, to reflect a greater degree of confidence in the proteomic and transcriptomic data than the epigenetic data, we weighted the epigenetic prizes by a factor of 0.33. The same methods for multi-omic integration were used for the ES and the cortical neuron networks.

To assess the robustness and specificity of our networks, we randomized the edges and prizes of our network solution, respectively, adding gaussian noise to the randomized edges. Nodes that were not previously assigned a prize (termed Steiner nodes) were filtered out if they appeared in fewer than 40 edge randomizations or more than 40 prize randomizations. Louvain community detection was then performed to create sub-network clusters. Within each sub-network, gProfiler2 version 0.2.0^148^ was used to identify significantly enriched biological processes via Gene Ontology Biological Process terms or REACTOME pathways. To identify the overlap between the KO and HD networks, we took the union of nodes and edges between the two networks and re-clustered the networks based on those nodes that were only found in the KO network, nodes found only in the HD network and the nodes found in both networks.

OmicsIntegrator requires selecting hyperparameters to visualize a genotype-specific network. These hyperparameters are the weights on the prizes (b), the network size (w) and the edge penalty (g). To choose a hyperparameter set, OmicsIntegrator was run on a set of parameters: β= ^3^, ω={ 1,3,6} and γ= {2,5,6}. The parameter choices were evaluated based on the mean node robustness and specificity, minimizing the KS statistic between prize and Steiner node degree, and the networks with high ratios of proteomic and transcriptomic prizes to epigenomic prizes. Networks were visualized using Cytoscape version 3.8.0.

## Resource availability

Materials and protocols will be distributed to researchers in a timely manner following publication. All data will be deposited in public data bases: RNAseq, ATACseq and ChIPseq in GEO, Protein in Massive.

This study did not generate new unique reagents. Further information and requests for resources and reagents should be directed to and will be fulfilled by the lead contact.

