## Supplemental Tables 1-4, 6 for "Intersecting impact of CAG repeat and Huntingtin knockout in stem cell-derived cortical neurons"

Table S1. Table of cell lines

| Line ID | CAG Repeat Alleles | CHDI ID | Line name | Assay |
|---|---|---|---|---|
| 20CAGn1* | 22/20 | CHDI-90001539 | RUES2 | Omics |
| 20CAGn2 | 20/20 | CHDI-90002887 | RUES2_20(20)CAG-cl66 | Omics |
| 20CAGn3 | 22/20 | CHDI-90001585 | RUES2_20(22)CAG-cl30 | Imaging |
| 20CAGn4 | 22/20 | CHDI-90002034 | RUES2_20(22)CAG-cl65 | Imaging |
| 56CAG | 56/22 | CHDI-90001590 | RUES2_56(22)CAG-cl23 | Omics |
| 72CAGn1 | 72/20 | CHDI-90002877 | RUES2_72(20)CAG-cl12 | Omics/ Imaging |
| 72CAGn2 | 72/20 | CHDI-90003146 | RUES2_72(20)CAG-cl2 | Imaging |
| 72CAGn3 | 72/20 | CHDI-90003147 | RUES2_72(20)CAG-cl9 | Imaging |
| HTTKO | None | CHDI-90002878 | RUES2_Htt_KO-cl8A | OMICS/ Imaging |

*Unedited parental line

Table S2. Number of immunocytochemistry experiments performed, and the number of cells analyzed across experiments for each marker in each cell line.

| **Line** | **No. of Experiments** | | | | | **Total No. of Cells** | | | | |
|---|---|---|---|---|---|---|---|---|---|---|
| | **BCL11B** | **DARPP32** | **Ki67** | **FOXG1** | **TBR1** | **BCL11B** | **DARPP32** | **Ki67** | **FOXG1** | **TBR1** |
| **20CAGn3** | **6** | **5** | **6** | **5** | **6** | 157503 | 123259 | 165932 | 173957 | 208537 |
| **20CAGn4** | **3** | **3** | **3** | **3** | **3** | 61967 | 66514 | 88350 | 43933 | 6648 |
| **72CAGn1(12)** | **6** | **5** | **6** | **6** | **6** | 161264 | 116440 | 127997 | 183529 | 175079 |
| **72CAGn2(2)** | **5** | **4** | **5** | **5** | **5** | 215935 | 161215 | 187407 | 194541 | 223921 |
| **72CAGn3(9)** | **3** | **3** | **4** | **3** | **3** | 101736 | 93563 | 113418 | 61064 | 95479 |
| **HTTKO** | **4** | **4** | **4** | **5** | **5** | 143054 | 127333 | 153447 | 142049 | 182798 |

Table S3: Number of immunocytochemistry experiments performed, and the number of cells analyzed across experiments for each marker by genotype after outliers were removed.

| **Line** | **No. of Experiments** | | | | | **Total No. of Cells** | | | | |
|---|---|---|---|---|---|---|---|---|---|---|
| | **BCL11B** | **DARPP32** | **Ki67** | **FOXG1** | **TBR1** | **BCL11B** | **DARPP32** | **Ki67** | **FOXG1** | **TBR1** |
| **20CAGn3/n4** | 5 | 4 | 5 | 5 | 4 | 136473 | 58726 | 136480 | 223881 | 98592 |
| **72CAGn1/2/3** | 7 | 6 | 7 | 7 | 4 | 369076 | 150327 | 255358 | 350443 | 194894 |
| **HTTKO** | 2 | 2 | 2 | 4 | 2 | 56836 | 59120 | 52029 | 70246 | 71805 |

Table S4: Number of differential Omics at the ES stage relative to control ES lines.

| Cell type | Group | Count | Assay | Direction |
|---|---|---|---|---|
| ES | 56CAG | 86 | Transcriptomics | Increased |
| ES | 56CAG | 112 | Transcriptomics | Decreased |
| ES | 72CAG | 66 | Transcriptomics | Increased |
| ES | 72CAG | 43 | Transcriptomics | Decreased |
| ES | HTT KO | 345 | Transcriptomics | Increased |
| ES | HTT KO | 208 | Transcriptomics | Decreased |
| ES | HTT KO | 0 | Proteomics | Increased |
| ES | HTT KO | 3 | Proteomics | Decreased |
| ES | 56CAG | 22 | Proteomics | Increased |
| ES | 56CAG | 98 | Proteomics | Decreased |
| ES | 72CAG | 11 | Proteomics | Increased |
| ES | 72CAG | 59 | Proteomics | Decreased |
| ES | HTT KO | 1359 | ATAC-seq | Increased |
| ES | HTT KO | 1602 | ATAC-seq | Decreased |
| ES | 56CAG | 200 | ATAC-seq | Increased |
| ES | 56CAG | 892 | ATAC-seq | Decreased |
| ES | 72CAG | 907 | ATAC-seq | Increased |
| ES | 72CAG | 862 | ATAC-seq | Decreased |
| ES | HTT KO | 19 | H3K4me1 ChIP-seq | Increased |
| ES | HTT KO | 68 | H3K4me1 ChIP-seq | Decreased |
| ES | 56CAG | 21 | H3K4me1 ChIP-seq | Increased |
| ES | 56CAG | 207 | H3K4me1 ChIP-seq | Decreased |
| ES | 72CAG | 8 | H3K4me1 ChIP-seq | Increased |
| ES | 72CAG | 36 | H3K4me1 ChIP-seq | Decreased |

Table S5: Differential genes between *HTT* KO and control, 56CAG repeat expansion and control, and 72CAG repeat expansion and control in RNA-seq, proteomics, ChIP-seq and ATAC-seq data from ES cells.

Additional excel file

Table S6: Number of differential Omics at the cortical neuron stage relative to control cell lines.

| **Cell Type** | **Group** | **Assay** | **Differential Counts** | **Genotype** |
|---|---|---|---|---|
| Cortical Neurons | Decreased | H3K4me1 ChIP-seq | 8249 | 56/22 |
| Cortical Neurons | Increased | H3K4me1 ChIP-seq | 4760 | 56/22 |
| Cortical Neurons | Decreased | ATAC-seq | 4604 | 56/22 |
| Cortical Neurons | Increased | ATAC-seq | 7072 | 56/22 |
| Cortical Neurons | Decreased | Proteomics | 341 | 56/22 |
| Cortical Neurons | Increased | Proteomics | 233 | 56/22 |
| Cortical Neurons | Decreased | Transcriptomics | 3864 | 56/22 |
| Cortical Neurons | Increased | Transcriptomics | 1382 | 56/22 |
| Cortical Neurons | Decreased | H3K4me1 ChIP-seq | 10728 | 72/20 |
| Cortical Neurons | Increased | H3K4me1 ChIP-seq | 5172 | 72/20 |
| Cortical Neurons | Decreased | ATAC-seq | 5953 | 72/20 |
| Cortical Neurons | Increased | ATAC-seq | 10634 | 72/20 |
| Cortical Neurons | Decreased | Proteomics | 1235 | 72/20 |
| Cortical Neurons | Increased | Proteomics | 999 | 72/20 |
| Cortical Neurons | Decreased | Transcriptomics | 4022 | 72/20 |
| Cortical Neurons | Increased | Transcriptomics | 1476 | 72/20 |
| Cortical Neurons | Decreased | H3K4me1 ChIP-seq | 4278 | KO |
| Cortical Neurons | Increased | H3K4me1 ChIP-seq | 3321 | KO |
| Cortical Neurons | Decreased | ATAC-seq | 3175 | KO |
| Cortical Neurons | Increased | ATAC-seq | 5828 | KO |
| Cortical Neurons | Decreased | Proteomics | 133 | KO |
| Cortical Neurons | Increased | Proteomics | 84 | KO |
| Cortical Neurons | Decreased | Transcriptomics | 739 | KO |
| Cortical Neurons | Increased | Transcriptomics | 425 | KO |

Table S7: Differential genes between *HTT* KO and control, 56CAG repeat expansion and control, and 72CAG repeat expansion and control in RNA-seq, proteomics, ChIP-seq, and ATAC-seq data from cortical neurons.

Additional excel file

Table S8: Nodes shared between both *HTT* KO and *HTT* CAG expanded repeat networks, as well as the fold changes of these genes relative to controls in RNA-seq, proteomics, ChIP-seq, and ATAC-seq data.

Additional excel file
