## Supplemental Figures 1-11 for "Intersecting impact of CAG repeat and Huntingtin knockout in stem cell-derived cortical neurons"

Figure S1

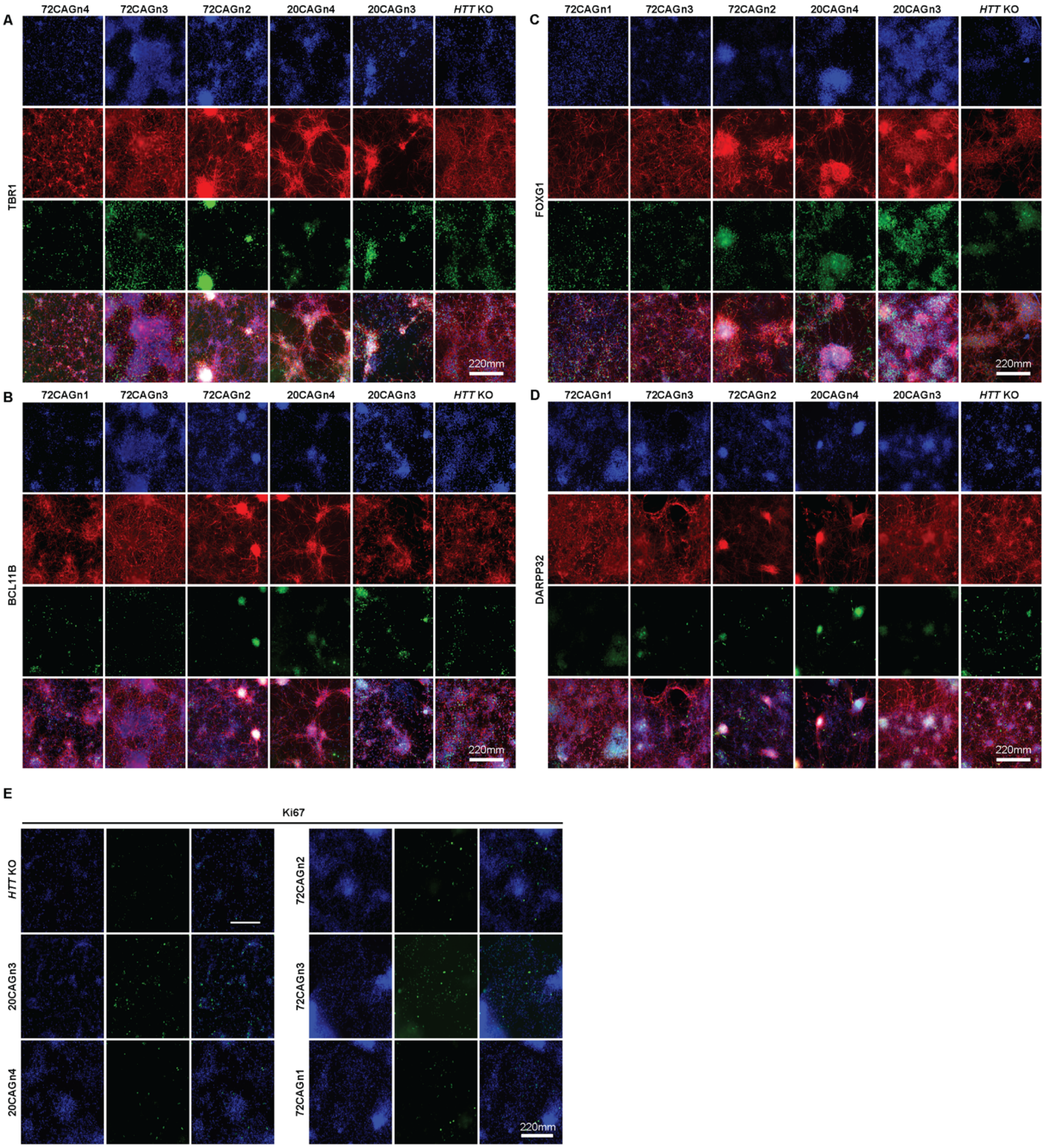


**Supplemental Figure 1: eCNs subjected to RM express cortical forebrain markers.** At differentiation day ~35 eCNs were fixed and stained with antibodies directed at various cell type specific markers from all clones (HTTKO, 20CAGn3, 20CAGn4, 72CAGn2, 72CAGn3, 72CAGn4). Representative immunofluorescence images showing expression of the neuronal forebrain markers **A**) TBR1 **B**) BCL11B

**C**) FOXG1. **D)** eCNs also patterned into cells that express the inhibitory GABAergic marker DARPP32. **E**) eCN cultures contained some proliferating cells as shown by the Ki67 stain. All cells stained with Hoescht (blue) nuclear marker. Panels **A-D** were also green with MAP2 (red). Scale bar = 220mm.

Figure S2

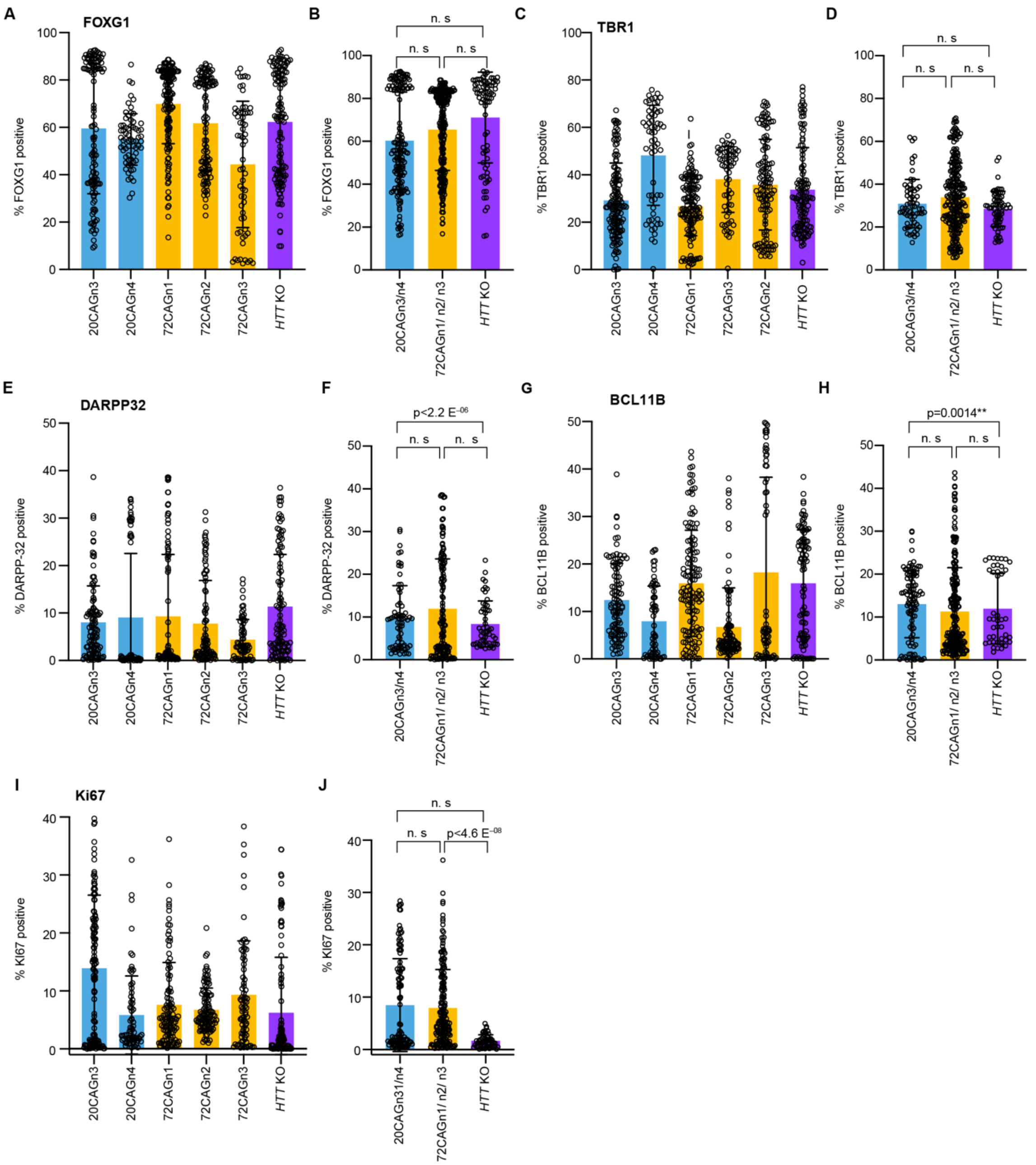


**Supplemental Figure 2: Quantification of day ~35 eCNs reveals a substantial proportion of neurons expressing typical forebrain markers.**

Day ~35 eCNs were fixed and stained with antibodies and Hoescht and imaged using RM. Images were subjected to a modified Cell Profiler pipeline as detailed in a previous study[1] to quantify the proportion of cell expressing each marker. We used a Linear Mixed Effects Model (LMM)[2,3] to quantify the percentage of cells positive for each antibody across each group. **A**, **C**, **E**, **G**, **I**) histograms showing the percentage of positive cells per image tile for each Hoechst- stained nuclei. **B**, **D**, **F, H, J**) histograms showing each group combined after outliers were removed as determined by the Cooks distance[4].

Figure S3

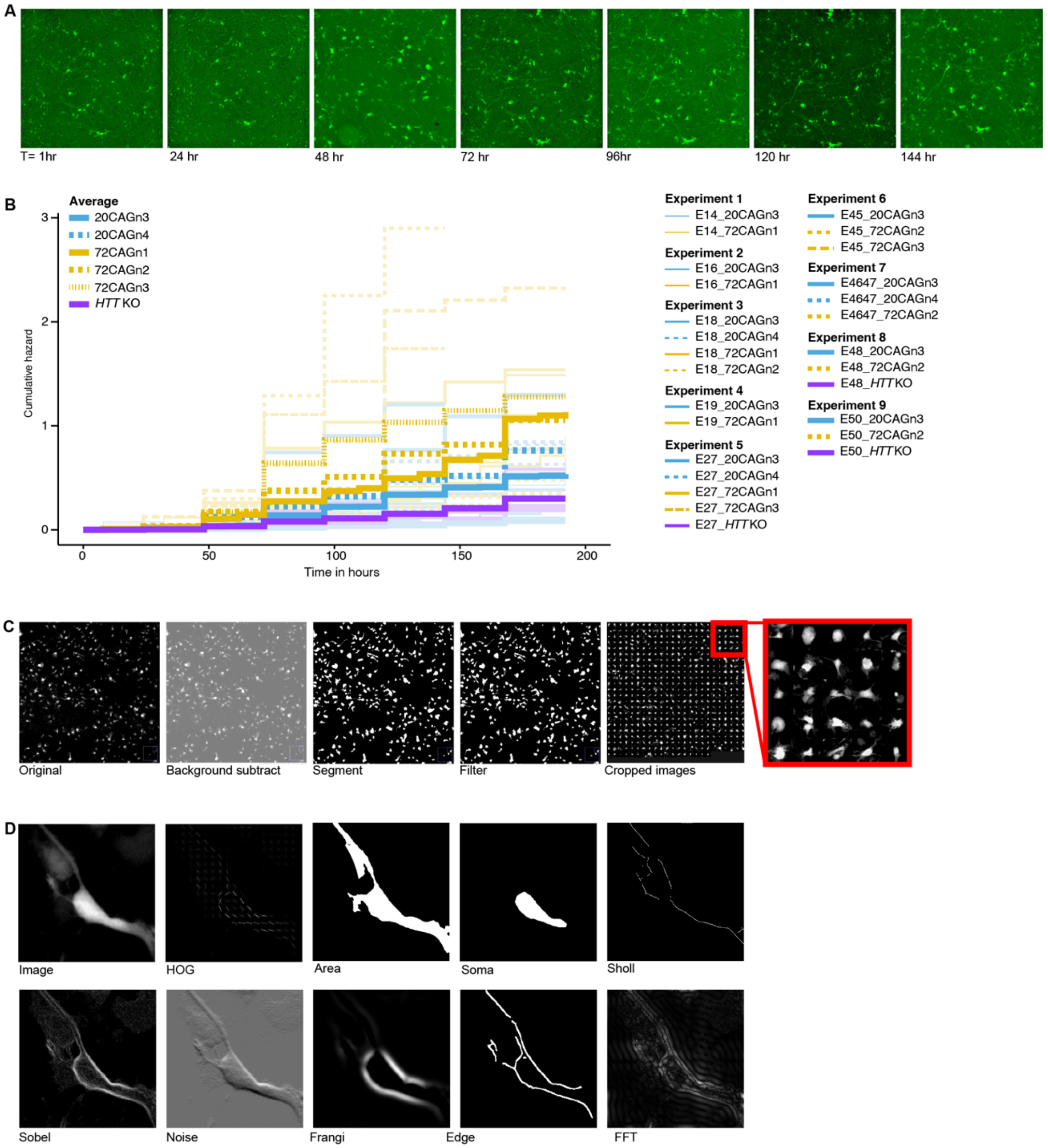


**Supplemental Figure 3: Image and feature analysis pipeline. A**) Example micrograph of live eCNs transduced with synapisn: EGFP imaged by RM every 24 hours. Scale bar = 65 mm. **B**) Cumulative risk of death curve plotted to show the variation across experiments. Each experiment is plotted by a different line thickness as delineated in the panels to the right. The average across all experiments are shown in bold. **C**) Example of image processing pipeline to generate single cell crops for ML. Each original image is acquired using RM. Images are subjected to our custom- built Galaxy pipeline that performs a background subtraction step, followed by segmentation based on the pixel intensity of each fluorophore. Next small or large objects are filtered to obtain single cells. These cells are then cropped to a 3 x3 image which is used for feature analysis. **D**) Example of feature maps shown for a single cropped neuron.

Figure S4

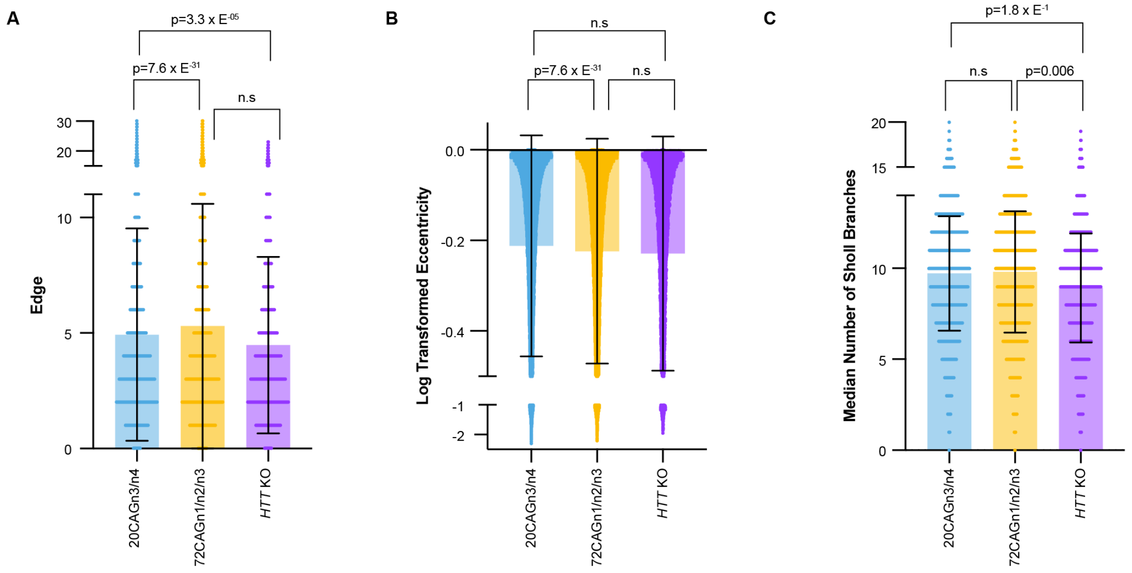


**Supplemental Figure 4: Comparison of eCN features at a single timepoint. A)** The perimeter of neuron including processes within each crop as captured by the Edge feature was larger in the 72CAG as compared to 20CAG and HTTKO. **B)** The shape or roundness of each cell as captured by Eccentricity showed that the 72CAG eCNs were rounder than the controls. **C)** The median number of processes emanating per soma captured by the Median Sholl feature was significantly different in the HTTKO eCNs as compared to control and 72CAG eCNs. Measurements were taken on images from differentiation day 28.

Figure S5

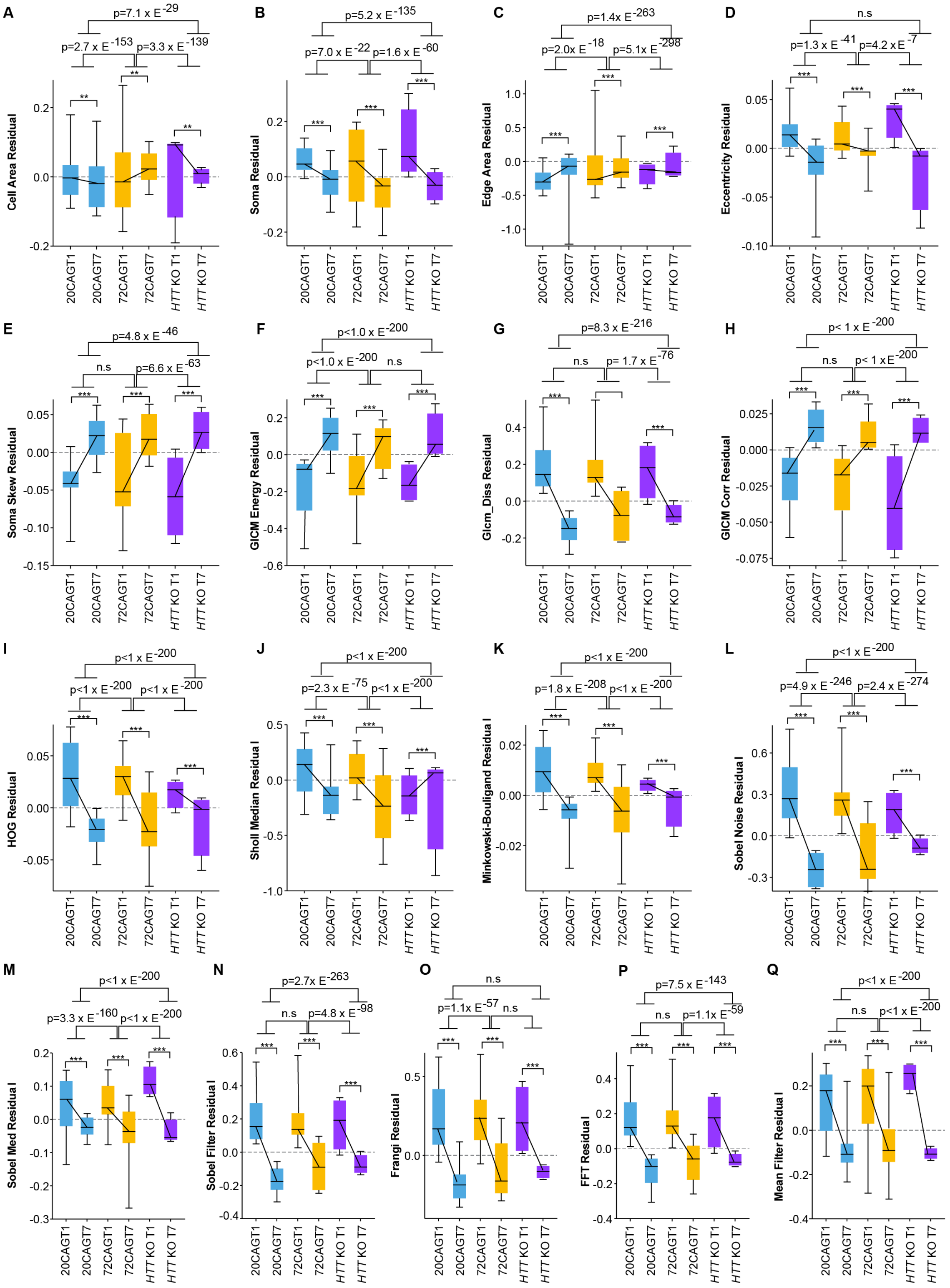

A
B
C
D
E
F
G
H
I
J
K
L
M
N
O
P
Q
Cell Area Residual
Soma Residual
Edge Area Residual
Eccentricity Residual
Soma Skew Residual
GlCM Energy Residual
Glcm_Diss Residual
GlCM Corr Residual
HOG Residual
Sholl Median Residual
Minkowski-Bouligand Residual
Sobel Noise Residual
Sobel Med Residual
Sobel Filter Residual
Frangi Residual
FFT Residual
Mean Filter Residual
20CAGT1
20CAGT7
72CAGT1
72CAGT7
HTT KO T1
HTT KO T7

**Supplemental Figure 5: Summary plots displaying feature changes over time. A)** The cell area feature captures the total cell area within each cropped image. The cell area was constant in the 20CAG and 72CAG eCNs over time, whereas cell area decreased significantly in the KO eCNs over time. Overall, the KO eCNs grew smaller over time by ~ 375% compared to the 72CAG or 20CAG CNs. **B).** The edge feature captures the perimeter of each neuron including processes within each crop. Overall, all eCNs displayed a reduced perimeter over time, suggesting decrease complexity over time. There was no difference in the rate of change of the perimeter between 20CAG and 72CAG eCNs, however, there was a decrease in the rate of change from the 20CAG and 72CAG to the KO eCNs by ~ 75%. This indicates that KO eCNs lose complexity faster than 20CAG and 72CAGs. **C)** The eccentricity (Edge Ecce) feature captures the shape or roundness of each cell. All eCNs became rounder over time, as the values drop closer to 0. The change in eccentricity was significantly different between 20CAG and 72CAG eCNs such that the controls became rounder 40% faster. Control and KO eCNs became round at the same rate. **D)** The grey level concurrence matrix correction (GLCM Corr) feature captures the texture of each neuron. All eCNs became smoother and less textured over time. The KO showed much faster rates of smoothing compared to the control (by ~130%) and the 72CAG by slightly less (by ~85%). **E)** The Haralick feature captures another type of texture based on intensity of each neuron but with different patterns than the GLCM. All lines displayed an increase in the Haralick feature over time. However, the 72CAG and KO eCNs increased more (~41% and 44% respectively) compared to the 20CAG eCNs. **F)** The histogram of gradients (HOG) feature captures complexity of structures and is similar to the edge feature, but captures pixels for the entire cell not just the edge. Over time, all eCNs became less complex according to this feature. There was no difference in the HOG feature for the 20CAG compared to the 72CAG eCNs. However, the KO eCNs displayed a ~70% lower rate of change compared to both 72CAG and 20CAG eCNs. **G)** The mean filter feature is another measure of pixel pattern complexity, and similarly decreased over time for all groups. The 72CAG eCNs decreased in this feature ~120% more than the 20CAG eCNs. **H)** The Minkowski-Bouligand (Mink Dim) feature is another complexity feature and also decreased over time in all groups. There was no difference between the control and 72 CAG eCNs however, both the control and 72CAG decreased faster than the KO eCNs by ~ 75%. **I)** The Sholl median (Sholl Med) features captures the median number of processes emanating from each soma. The number of processes decreased over time in the control and 72CAG eCNs, whereas in the KO eCNs the number of processes increased. **J)** The Sobel med feature is another edge detection feature. This feature was static in the control eCNs while it decreased in the 72CAG and KO. This feature decreased more significantly in the 72CAG eCNs compared to controls by ~250%. **K)** The Sobel noise is another version of an edge detection feature. This feature decreased across all eCNs over time. There was no difference from control to the 72CAG eCNs, however the feature decreased more slowly in KO eCNs than the 72CAG and control eCNs by ~ 30-40% respectively. **L)** The soma area feature (soma) captures the size of just the soma. There is no change in the control and 72CAG eCNs over time. However, the KO eCNs are decreasing is size to the controls and 72CAG by ~500 and 300% respectively. **M)** The soma med feature captures the median pixel value intensity across the soma. This feature decreased over time across all lines, but faster in the 72CAG eCNs compared to controls by ~36%. **N)** The soma skew feature measures the symmetry gradient of pixels across the soma. The soma skew feature values increased over time in all lines, but faster in the 72CAG eCNs are increasing faster than the controls by ~22%.

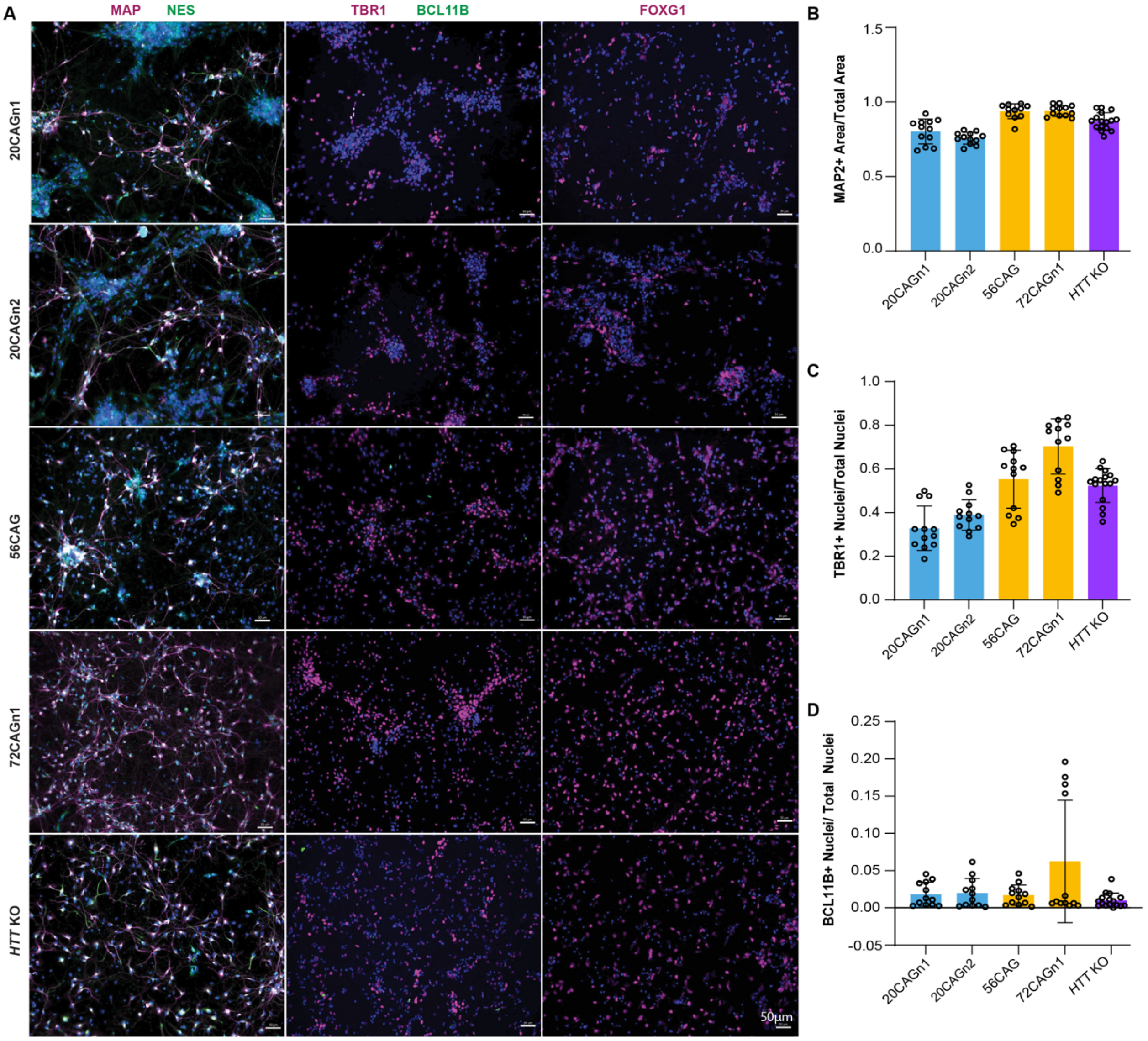


**Supplemental Figure 6: eCNs for OMIC analysis display similar cell type characterization. A).** Representative immunofluorescence images of omics studies eCN d35 differentiations for all cell lines (20CAGn1, 20CAGn2, 56CAG, 72C.AGn1, and HTTKO) showing high expression of mature neuronal marker MAP2 (magenta) and low expression of immature neuronal marker NES (green), as well as widespread expression of forebrain marker FOXG1 (magenta). Cells display very low expression of post mitotic neuronal marker BCL-11B (green) but high expression of cortical layer marker TBR1 (magenta). All cells stained with Hoescht (Blue) nuclear marker. Scale bar= 50um. **B).** Bar graph showing the fraction MAP2+ positive cell bodies by area per cell line (n=12-16 images per cell line across 3 (4 for HTTKO) differentiations). **C).** Bar graph showing the fraction TBR1 positive nuclei normalized to total nuclei (n=12-16 images per cell line across 3 (4 for HTTKO) differentiations). **D).** Bar graph showing the fraction CTIP2 positive nuclei normalized to total nuclei (n=12-16 images per cell line across 3 (4 for HTTKO) differentiations). Error bars represent standard deviation.

Figure S7

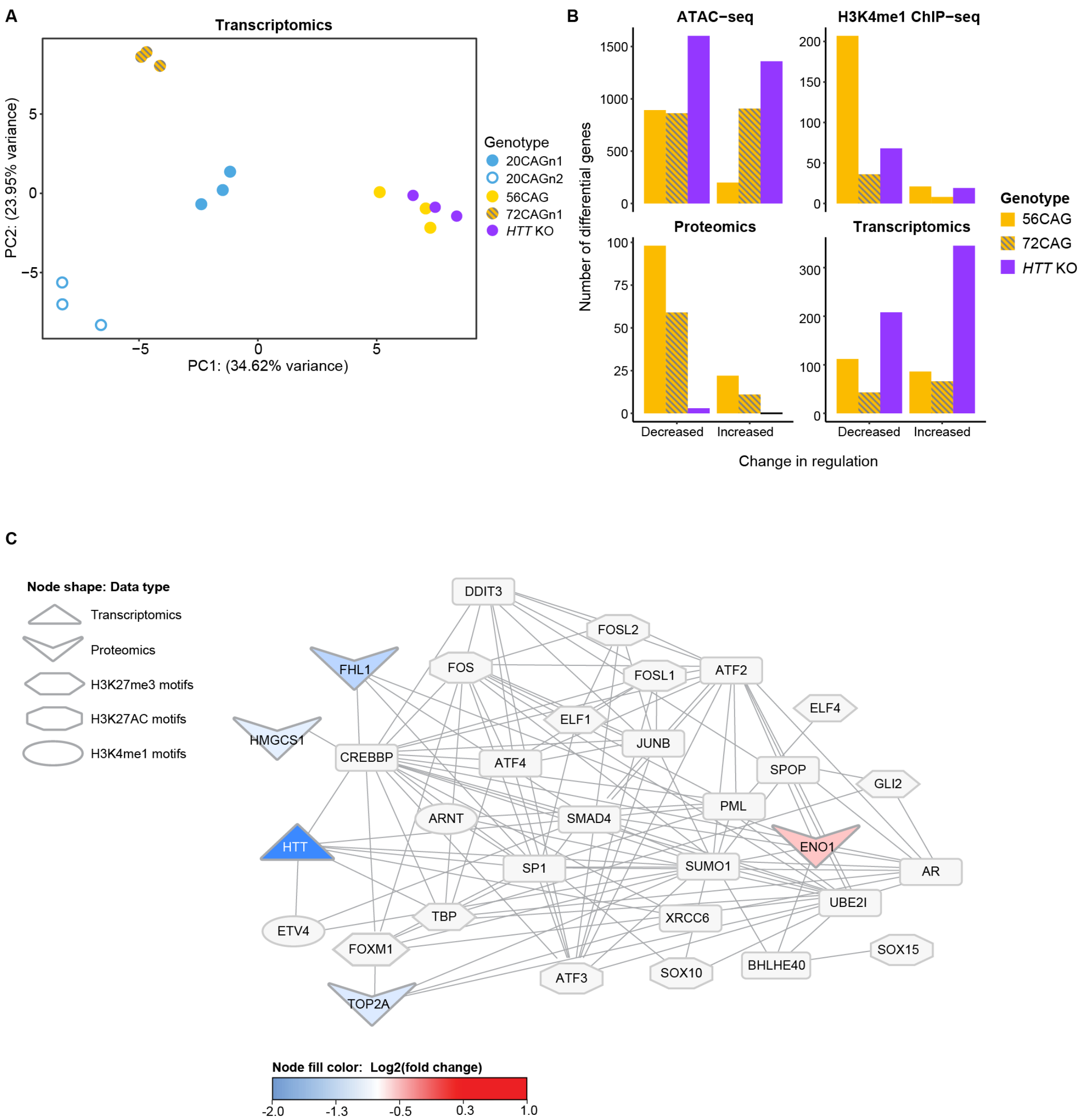


**Supplemental Figure 7: Pluripotent stage omics analysis. A)** PCA of pluripotent cell transcriptomics shows clustering by cell line across multiple replicates but not by HD status. Parentheses indicate percentage of variance explained by principal components 1 and 2. **B)** Number of differential genes comparing HTT KO to control, 56CAG to control and 72CAG to control in ES cells measured by ATAC-seq, H3K1 ChIP-seq, proteomics and transcriptomics. 56CAG is indicated in the gold bars, 72CAG is indicated in the gold bars and *HTT*KO is indicated in the purple bar. For the ATAC-seq and H3K4me1 ChIP-seq, we defined differential genes as genes for which there was at least one significant peak at FDR<0.1. C) A subnetwork from the integrative analysis of *HTT* KO multi-omic data in ES cells shows interactions between the down-regulated HTT protein with known its known interactor CREBBP, which in turn interacts with its known downstream effector FOS. Other interacting nodes such as the down-regulated protein TOP2A and FOXM1 are involved in DNA repair maintenance.

Figure S8

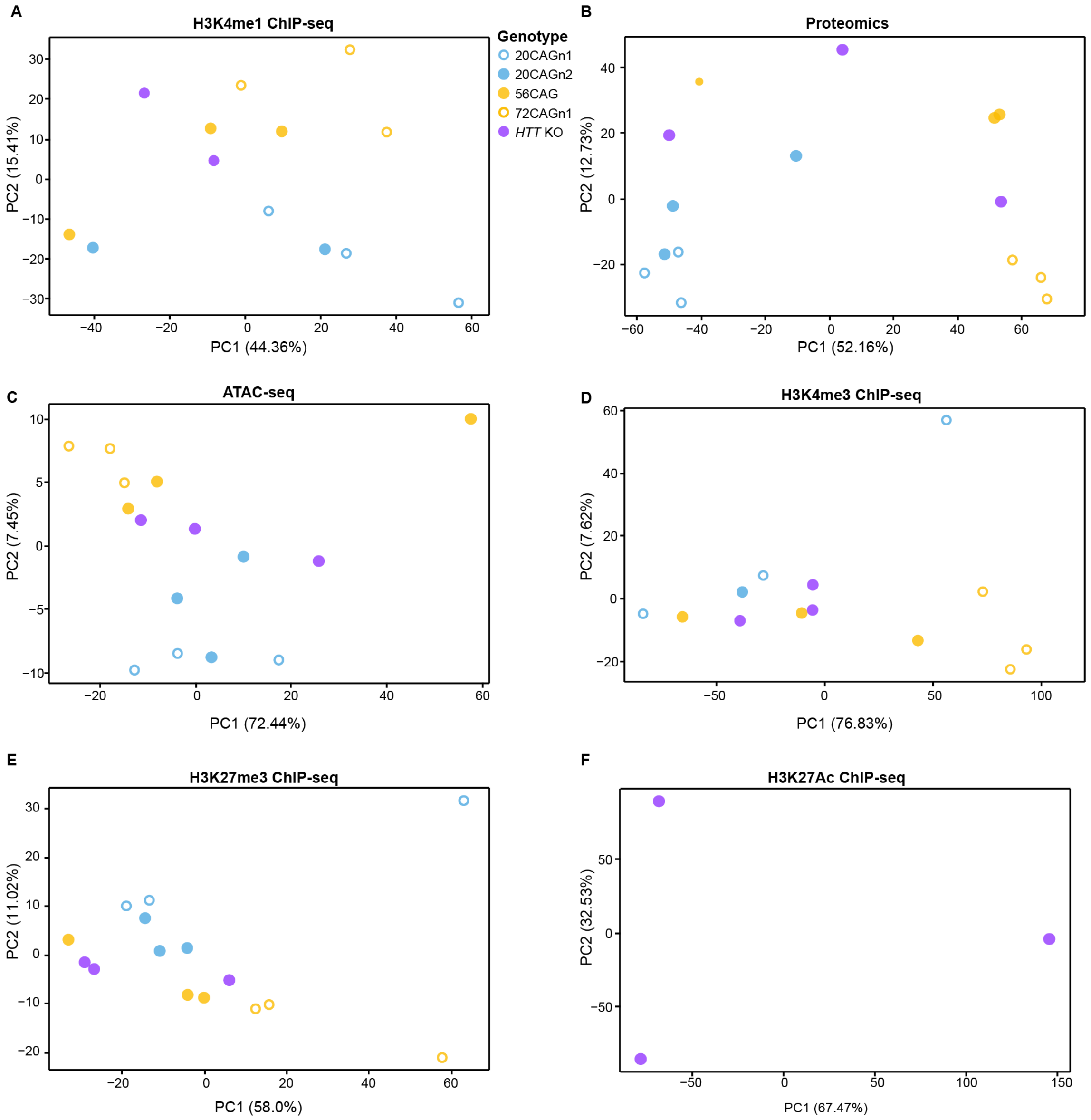


**Supplemental Figure 8: Primary Omics assays in ESC-derived cortical neurons**. A) PCA of H3K4me1 ChIP-seq shows partial separation by HD status along principal component 2. One replicate (20CAGn1, replicate 2) in the control in the H3K4me1 data was removed and excluded from downstream analysis. B) PCA of proteomics separates some cell lines by HD status along principal component 1. C) PCA of ATAC-seq separates some cell lines by HD status along principal component 2. One replicate (56CAG, replicate 3) in the ATAC-seq data was removed from the PCA for failing quality control benchmarks due to low read counts and was excluded from downstream analysis. D) PCA of H3K4me3 ChIP-seq separates some cell lines by HD status along principal component 1. E) PCA of H3K27me3 ChIP-seq separates some cell lines by HD status along principal component 2. F) PCA of H3K27Ac ChIP-seq separates some cell lines by replicate ID rather than genotype. Parentheses in panels B-D and F-G indicate percentage of variance explained by principal components 1 and 2.

Figure S9

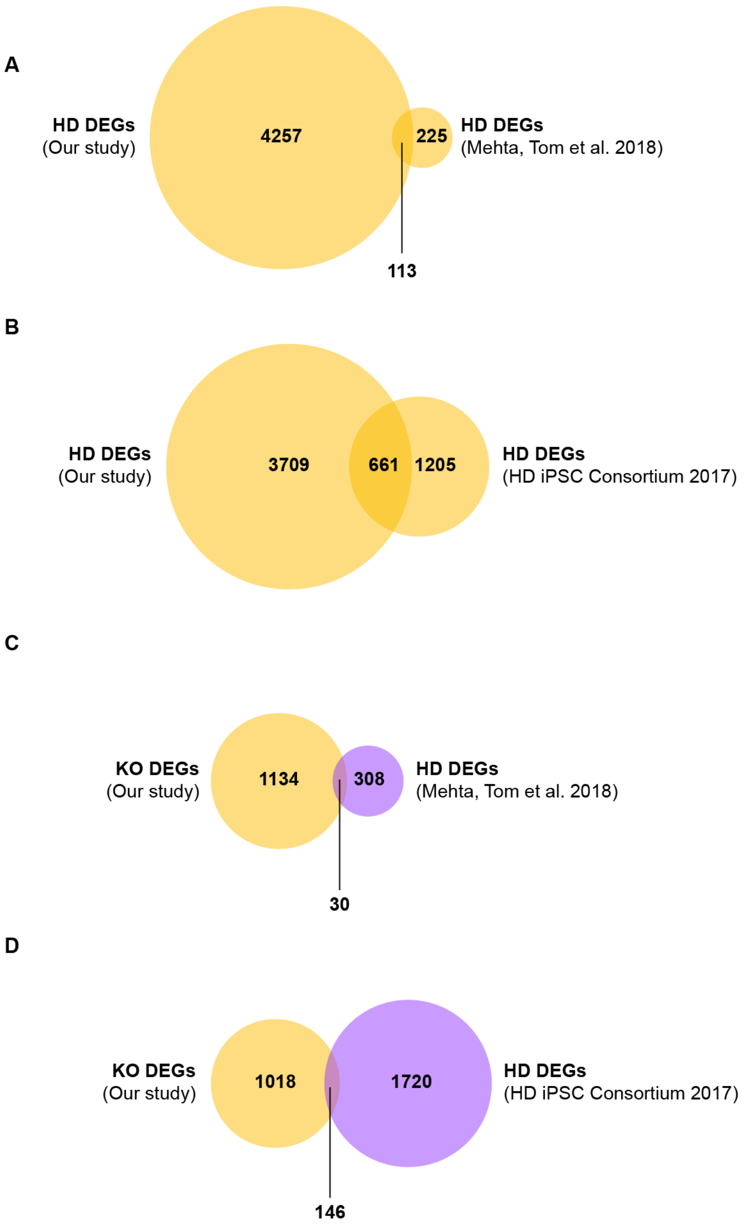


**Supplemental Figure 9: Comparison of transcriptional changes with previously published studies.** A) The differentially expressed genes between HD case and control transcriptomics significantly overlap with the differentially expressed genes between HD and control in from a previous study using iPSC-derived cortical neurons (hypergeometric p-value=$2.0*10^{-6}$). B) The differentially expressed genes between HD case and control transcriptomics significantly overlap with previously published differentially expressed genes between HD and control from iPSC-derived striatal neurons (hypergeometric p-value<$10^{-16}$). C) The differentially expressed genes between *HTT*KO and control have marginally significant overlap with the differentially expressed genes between HD and control in previously published iPSC-derived cortical neurons (p=0.02). D) The differentially expressed genes between *HTT*KO and control significantly overlap with differentially expressed genes between HD and control from a previously published study of iPSC-derived striatal neurons (p=$3.8*10^{-4}$).

Figure S10

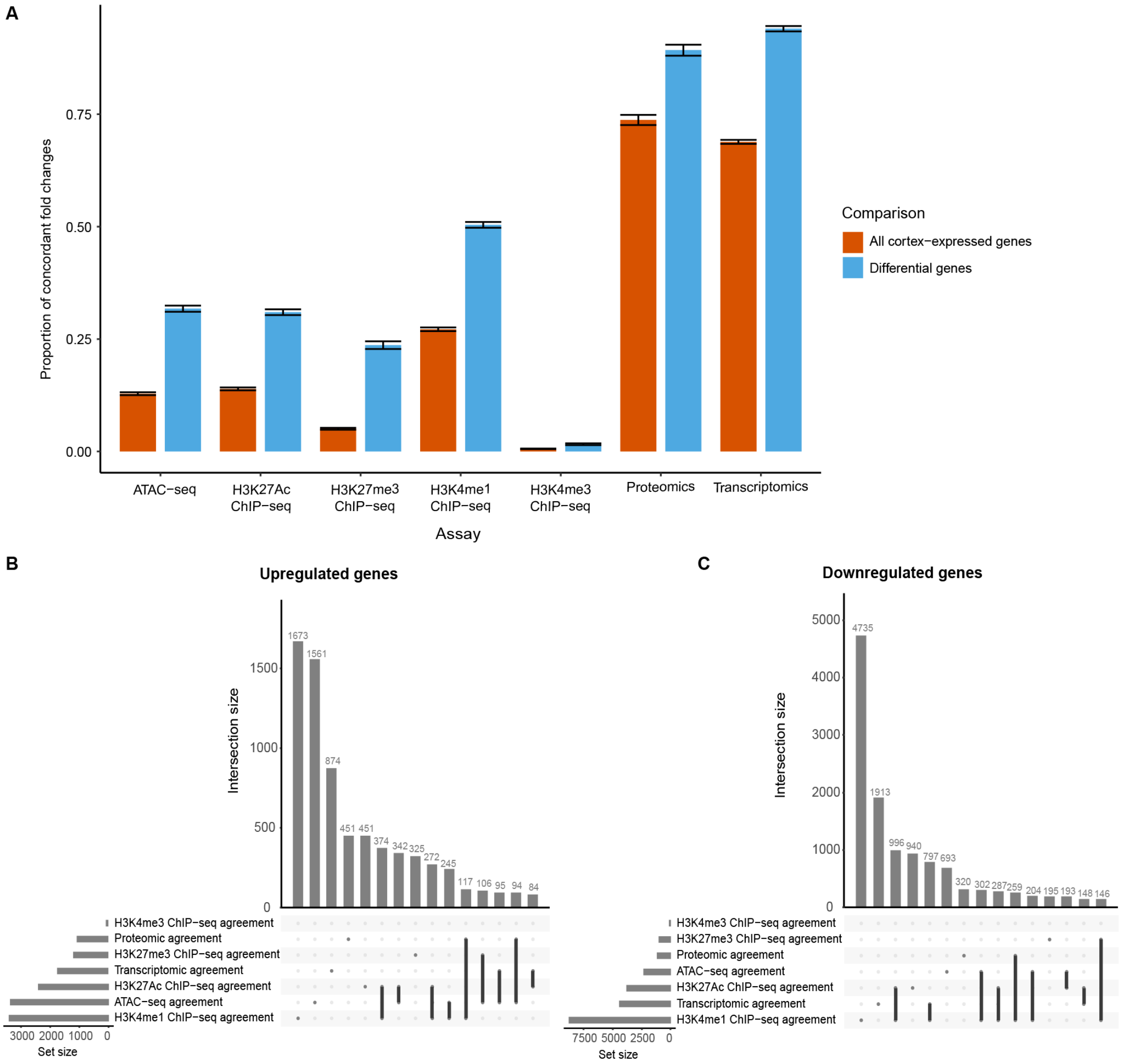


**Supplemental Figure 10: Summary of concordant changes analysis. A):** Differentially expressed genes between HD and control or *HTT* and control are enriched for genes with the same fold change across genotypes in each assay. UpSet plots of upregulated and downregulated genes show that genes that are concordant in fold change across multiple assays are among the largest intersection sets for B) upregulated genes and C) downregulated genes.

Figure S11

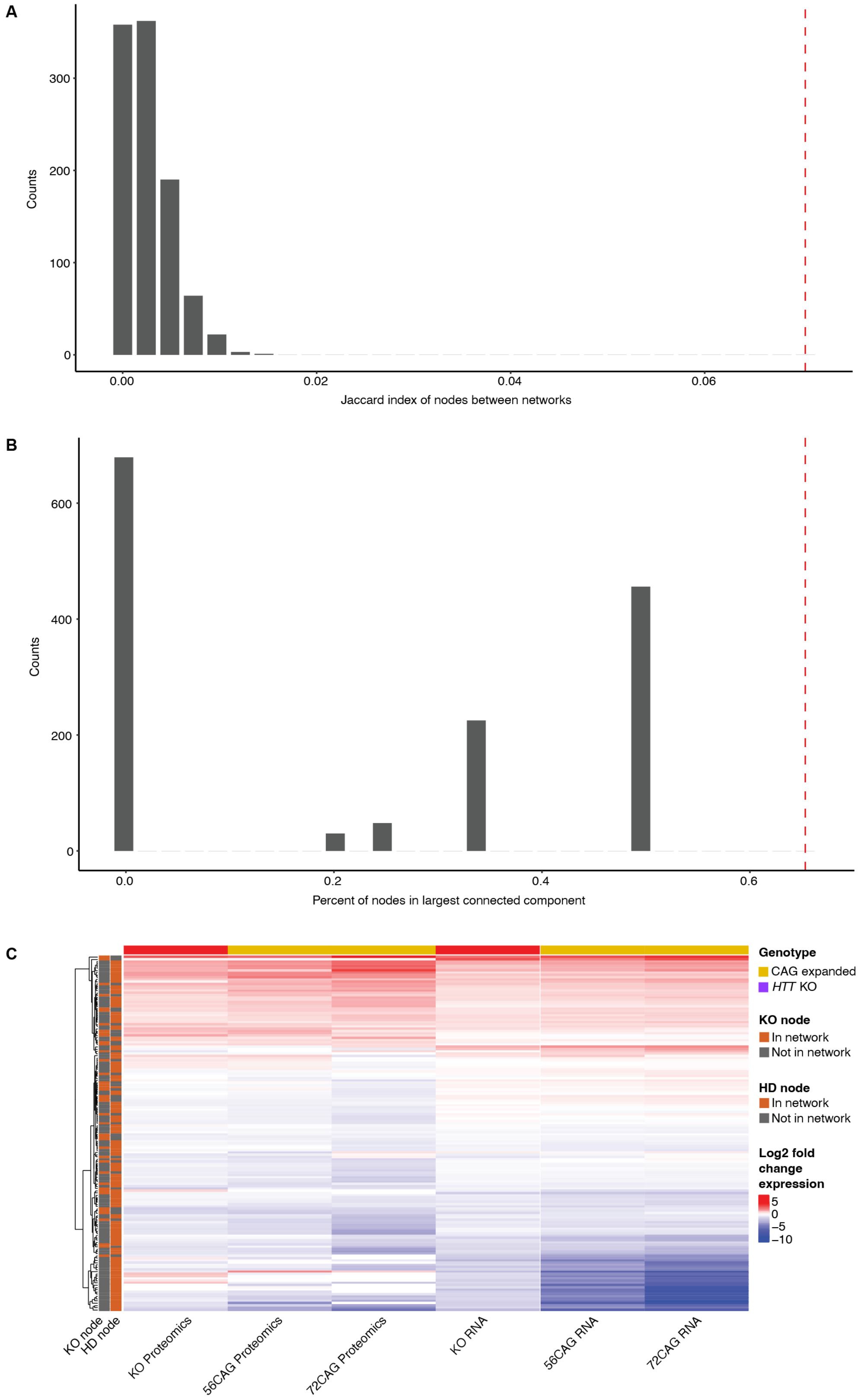
A
Counts
300
200
100
0
0.00
0.02
0.04
0.06
Jaccard index of nodes between networks
B
Counts
600
400
200
0
0.0
0.2
0.4
0.6
Percent of nodes in largest connected component
C
Genotype
CAG expanded
HTT KO
KO node
In network
Not in network
HD node
In network
Not in network
Log2 fold change expression
5
0
−5
−10
KO node
HD node
KO Proteomics
56CAG Proteomics
72CAG Proteomics
KO RNA
56CAG RNA
72CAG RNA

**Supplemental Figure 11: Degree of overlap between CAG expanded and KO networks A)** The number of intersecting nodes between the *HTT* KO and HD networks are greater than expected by chance after measuring the Jaccard index of 1000 pairs of randomized networks. The Jaccard index between the *HTT* KO and HD networks is indicated by the dashed, vertical red line. **B)** The size of the largest connected component at the intersection of the *HTT* KO and HD network is larger than the largest connected component of any of 1000 other randomized networks. The percentage of nodes in the largest connected component is indicated by the dashed, vertical red line. **C)** Heatmap of the fold change between case and control in proteomics and transcriptomics for nodes that appear in the HTT KO network or the HD network. Many of these nodes have similar fold change directions across assays.

1. Wang B, Vartak R, Zaltsman Y, Naing ZZC, Hennick KM, Polacco BJ, Bashir A, Eckhardt M, Bouhaddou M, Xu J, Sun N, Lasser MC, Zhou Y, McKetney J, Guiley KZ, Chan U, Kaye JA, Chadha N, Cakir M, Gordon M, Khare P, Drake S, Drury V, Burke DF, Gonzalez S, Alkhairy S, Thomas R, Lam S, Morris M, Bader E, Seyler M, Baum T, Krasnoff R, Wang S, Pham P, Arbalaez J, Pratt D, Chag S, Mahmood N, Rolland T, Bourgeron T, Finkbeiner S, Swaney DL, Bandyopadhay S, Ideker T, Beltrao P, Willsey HR, Obernier K, Nowakowski TJ, Huttenhain R, State MW, Willsey AJ, Krogan NJ: A foundational atlas of autism protein interactions reveals molecular convergence. bioRxiv 2024. PMC10705567.
2. Kuznetsova A, Brockhoff PB, Christensen RHB: lmerTest package: Tests in linear mixed effects models. J Stat Softw 2017, 82:1–26.
3. R Core Team: R: A language and environment for statistical computing. Edited by Vienna, Austria, R Foundation for Statistical Computing. Available from: https://www.R-project.org/, 2012, p. PMCID.
4. Cook RD: Influential Observations in Linear Regression. Journal of the American Statistical Association 1979, 74.
